# An RNA Damage Response Network Mediates the Lethality of 5-FU in Clinically Relevant Tumor Types

**DOI:** 10.1101/2023.04.28.538590

**Authors:** Jung-Kuei Chen, Karl A. Merrick, Yi Wen Kong, Anita Izrael-Tomasevic, George Eng, Erika D. Handly, Jesse C. Patterson, Ian G. Cannell, Lucia Suarez-Lopez, Aaron M. Hosios, Anh Dinh, Donald S. Kirkpatrick, Kebing Yu, Christopher M. Rose, Jonathan M. Hernandez, Haeun Hwangbo, Adam C. Palmer, Matthew G. Vander Heiden, Ömer H. Yilmaz, Michael B. Yaffe

## Abstract

5-fluorouracil (5-FU) is a successful and broadly used anti-cancer therapeutic. A major mechanism of action of 5-FU is thought to be through thymidylate synthase (TYMS) inhibition resulting in dTTP depletion and activation of the DNA damage response. This suggests that 5-FU should synergize with other DNA damaging agents. However, we found that combinations of 5-FU and oxaliplatin or irinotecan failed to display any evidence of synergy in clinical trials, and resulted in sub-additive killing in a panel of colorectal cancer (CRC) cell lines. In seeking to understand this antagonism, we unexpectedly found that an RNA damage response during ribosome biogenesis dominates the drug’s efficacy in tumor types for which 5-FU shows clinical benefit. 5-FU has an inherent bias for RNA incorporation, and blocking this greatly reduced drug-induced lethality, indicating that accumulation of damaged RNA is more deleterious than the lack of new RNA synthesis. Using 5-FU metabolites that specifically incorporate into either RNA or DNA revealed that CRC cell lines and patient-derived colorectal cancer organoids are inherently more sensitive to RNA damage. This difference held true in cell lines from other tissues in which 5-FU has shown clinical utility, whereas cell lines from tumor tissues that lack clinical 5-FU responsiveness typically showed greater sensitivity to the drug’s DNA damage effects. Analysis of changes in the phosphoproteome and ubiquitinome shows RNA damage triggers the selective ubiquitination of multiple ribosomal proteins leading to autophagy-dependent rRNA catabolism and proteasome-dependent degradation of ubiquitinated ribosome proteins. Further, RNA damage response to 5-FU is selectively enhanced by compounds that promote ribosome biogenesis, such as KDM2A inhibitors. These results demonstrate the presence of a strong RNA damage response linked to apoptotic cell death, with clear utility of combinatorially targeting this response in cancer therapy.

## Introduction

DNA damaging chemotherapeutic drugs are the mainstay of medical treatment for many tumor types, including colorectal cancer. These drugs typically cause genotoxic stress by directly modifying DNA bases, cross-linking DNA strands, or inhibiting enzymes required for genome replication (Pearl et al., 2015; Reuvers et al., 2020). These genomic alterations activate the DNA damage response (DDR), a broad signaling network that arrests the cell division cycle, recruits DNA repair machinery to the lesions, and controls the onset of cell cycle re-entry, senescence or programmed cell death (Ciccia and Elledge, 2010; Matt and Hofmann, 2016). Despite intensive efforts to understand the determinants of response to chemotherapeutics, it is frequently difficult to predict which patients will respond to which DNA damaging drugs, how to best combine different DNA damaging agents with each other, or whether small molecule inhibitors of the DNA damage response will affect patient outcomes.

Most combination chemotherapy regimens currently used in clinical practice originated from empiric observations and clinical trials that were often designed based on non-overlapping toxicities (Pritchard et al., 2012). Among these, the combination of oxaliplatin and/or irinotecan with 5-fluorouracil (5- FU) and leucovorin (denoted FOLFOX and FOLFIRI, respectively) has shown modest efficacy in the treatment of patients with advanced stage gastrointestinal malignancies, particularly pancreatic and colorectal cancer, that exceeds that observed with 5-FU plus leucovorin alone. (de Gramont et al., 2000; Saltz et al., 2000), While all three chemotherapeutic agents cause DNA damage, each has a distinct mechanism of action suggesting the possibility of synergy. Platinum agents function by inducing intrastrand and interstrand DNA crosslinks between purine bases (Dasari and Tchounwou, 2014; Deans and West, 2011). Irinotecan is a camptothecin analog that forms a ternary complex with topoisomerase I and DNA, resulting in single- and double-strand DNA breaks. 5-FU, which is present in both the FOLFOX and FOLFIRI regimens, has multiple activities. Following cell entry, 5-FU is converted to several active metabolites, including 5-fluorodeoxyuridine monophosphate (5-FdUMP), 5-fluorodeoxyuridine triphosphate (5-FdUTP), and 5-fluorouridine triphosphate (5-FUTP). 5-FdUMP potently inhibits substrate binding of dUMP to TYMS, resulting in depletion of dTTP pools, impaired DNA synthesis and replication fork stalling. The 5-FdUTP metabolite itself can be directly incorporated into DNA, contributing to DNA damage that requires base excision repair (Thorn et al., 2011). In addition to the DNA damaging metabolites described above, 5-FU treatment also leads to the incorporation of the 5-FUTP into RNA, which affects the processing and maturation of multiple RNA species (elegantly reviewed in (Longley et al., 2003). The extent to which these damaged RNAs contribute to 5-FU cytotoxicity in mammalian tumors, or if they are cleared from the RNA pool while not being detected or having negligible cytotoxic effects remains a topic of ongoing and active debate. Recent papers continue to emphasize important roles for 5-FU-mediated DNA damage (Ludikhuize et al., 2022), and the fact that a current clinical standard of care for hepatic metastases from colorectal cancer is direct administration of the deoxynucleoside 5-fluorodeoxyuridine (Floxuridine) into the liver through a hepatic artery infusion pump (Doussot et al., 2015), reflects the general belief that major clinical benefits of 5-FU derive from its DNA-damaging deoxy metabolites.

Despite the distinct mechanisms of action of oxaliplatin and irinotecan, their clinical efficacy is indistinguishable when either is combined with 5-FU (Colucci et al., 2005). One possible explanation for these beneficial clinical effects seen with irinotecan/5-FU and oxaliplatin/5-FU combination therapy could be that 5-FU disruption of deoxynucleotide pools further compromises the ability of the cell to repair of irinotecan- or oxaliplatin-induced DNA lesions, or that concomitant incorporation of 5-FdUTP into DNA enhances the lethality of the DNA damage induced by DNA crosslinking or double strand break formation. We therefore explored interactions between these DNA damaging agents to define the contexts in which they are optimally synergistic and to potentially identify novel targeted agents to enhance the response. We made the unexpected observation that the primary mechanism by which 5-FU kills colorectal and cancer cell lines, as well as cell lines from other clinically relevant indications is through an RNA damage-induced apoptotic pathway. We specifically demonstrate that 5-FU induced cytotoxicity results from disruption of ribosome biogenesis, resulting in nucleolar restructuring, lysosomal degradation of ribosomal RNA, and proteasomal degradation of ubiquitinated ribosomal protein subunits, leading to apoptotic cell death in both p53 wild-type and mutant cell lines. We show that modulating the rate of ribosome biogenesis using nutrients, small molecule inhibitors, or genetic manipulation can modulate the sensitivity of CRC cells to 5-FU-induced cell death, indicating the potential to further target ribosome biogenesis pathways for improved anti-cancer treatment options.

## RESULTS

### 5-fluorouracil does not enhance the efficacy of oxaliplatin or irinotecan in killing CRC cells, nor display synergy in human clinical trials

To examine the cell killing efficacy of oxaliplatin and irinotecan, two standard DNA damaging chemotherapeutic agents used in the treatment of colorectal cancer, we screened 11 well-established colorectal cancer cell lines for their sensitivity to these drugs, as well as to 5-FU (Figure 1A and B). Relative drug sensitivities between the different cell lines were compared by integrating the area under the dose-survival curves (Figure 1B), revealing a wide range of responses to each of the different drugs. Some cell lines were uniformly sensitive to all three drugs (i.e. SW48, HCT116, LoVo), others were moderately resistant to all of the drugs (i.e. HT- 29, GP5d), while still others showed strong a varied resistance to one or two of the agents, likely recapitulating the clinical response seen in a heterogeneous tumor population.

**Figure 1:**
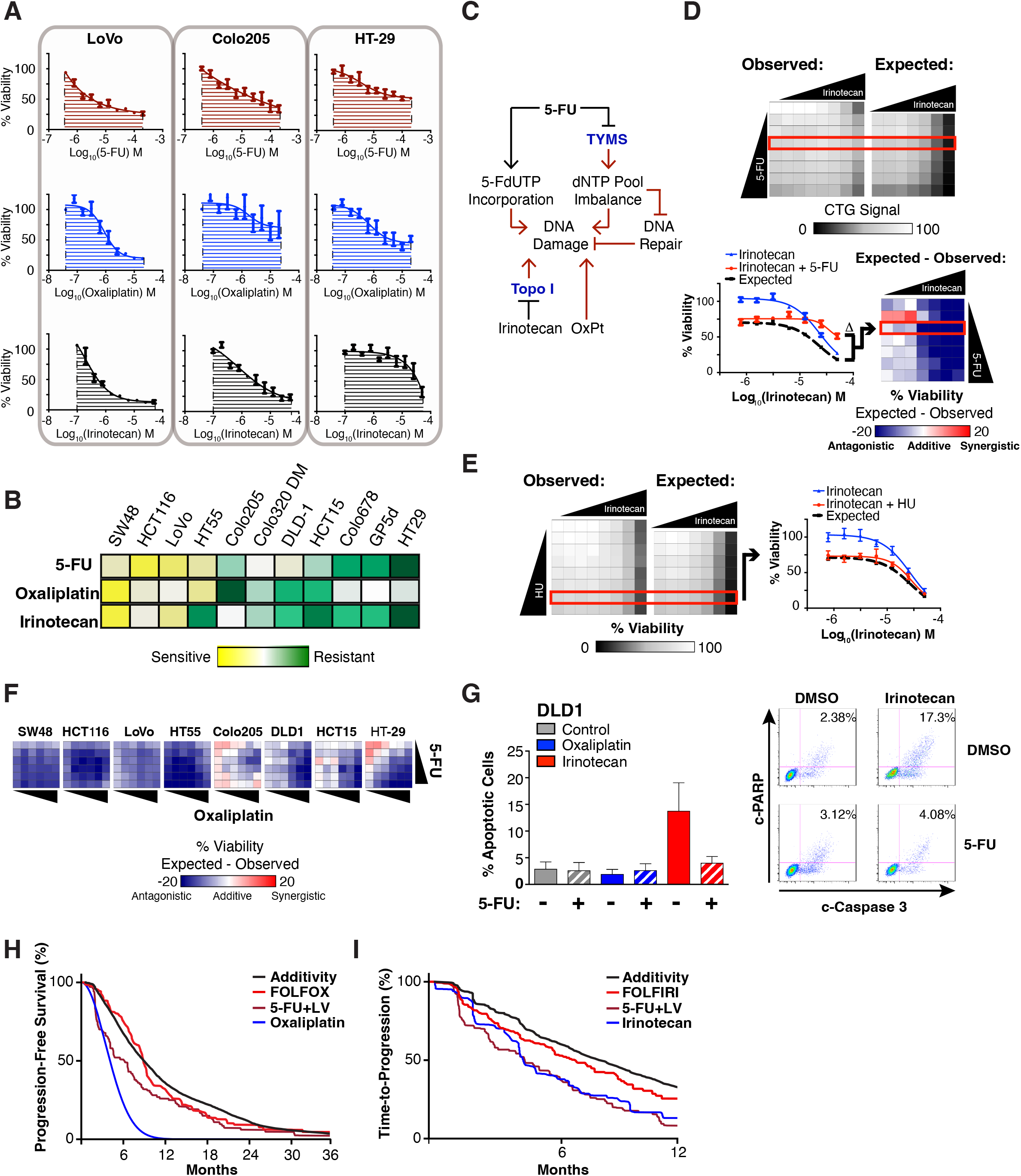
Sub-additivity and antagonism of tumor cell killing in clinically relevant 5-FU drug combinations. A) Sensitivity of LoVo, Colo205, and HT-29 CRC cells to 5-FU, oxaliplatin, and irinotecan was determined using Resazurin cell viability assay after 72 h of drug treatment. The area under the curve (AUC) used to summarize the drug response in panel B is highlighted. B) Area under the curve data for each drug response was determined as in A) reveals distinct sensitivities of CRC cell lines to 5-FU, oxaliplatin, and irinotecan. C) 5-FU treatment impacts the DDR in multiple ways to potentially alter response to oxaliplatin and irinotecan. D) HT-29 cells were treated with a dose-response matrix of 5-FU and irinotecan (top doses of 200 μM and 50 μM, respectively), and cell viability was measured after 72 hours by CellTiter-Glo (Observed). The effect of each drug individually was used to calculate the expected viability of the drug combinations based on a Bliss Independence model (Expected), and the effect of the drug combination was determined by comparing these values. Negative numbers indicate antagonism or sub-additivity, 0 indicates an additive effect, and positive numbers indicate synergy. E) HT-29 cells were treated with a dose-response matrix of hydroxyurea (HU) and irinotecan (top doses of 4 mM and 50 μM, respectively). Observed and expected viability from the combinations was determined as in panel F. F) Dose-response matrices for the indicated cell lines were generated with 5-FU and oxaliplatin (top doses of 200 μM and 20 μM, respectively), and viability was measured using Resazurin cell viability assay after 72 h of drug treatment. The effect of the drug combinations was determined as in panel D. G) DLD1 cells were treated with oxaliplatin (5 μM) or irinotecan (6.25 μM) +/- 50 μM 5- FU for 48 hours, and the induction of apoptosis was measured by flow cytometry. H) 5-FU/LV and irinotecan have sub-additive effects on time to progression (hazard ratio for observed versus expected efficacy = 1.25, 95% CI [1.03, 1.51], Cox proportional hazard model). Note that HR=1 does not mean no efficacy, it means as effective as expected, Cox proportional hazard model. I) 5-FU/LV and oxaliplatin have additive effects on Progression-free survival (hazard ratio for observed versus expected efficacy = 1.08, 95% CI [0.86, 1.36], Cox proportional hazard model).

Oxaliplatin and irinotecan cause distinct types of DNA damage that utilize different molecular mechanisms for repair. Intra-strand DNA crosslinks formed by oxaliplatin are typically excised by the nucleotide excision repair (NER) pathway or bypassed by the translesion synthesis (TLS) pathway, while inter-strand crosslinks are repaired primarily by the Fanconi pathway and/or homologous recombination. Single- and double-strand DNA breaks generated by irinotecan inhibition of topoisomerase I are repaired primarily by single strand annealing, homologous recombination, or non-homologous end joining (Pommier, 2006). Despite these differences, all of these repair mechanisms require the action of DNA polymerases to fill in gaps generated during the excision and repair process itself and likely require maintenance of dNTP pools for efficient repair (Chabes et al., 2003). Thus, depletion of the free dNTP pool as a consequence of 5-FU-mediated inhibition of TYMS, along with inappropriate incorporation of 5-FdUTP into DNA would be expected to synergistically enhance the cell death caused by oxaliplatin or irinotecan at one or more doses (Figure 1C). We therefore performed detailed dose-response measurements for cell survival in response to combination treatment with irinotecan and 5-FU or oxaliplatin and 5-FU. Surprisingly, the effect of combining 5-FU with oxaliplatin or irinotecan resulted in sub-additive loss of viability in nearly all cell lines examined (Figure 1D and F, Supplemental Figure 1A). Conversely, we observed clear additivity between another nucleotide depleting drug, hydroxyurea, and irinotecan, suggesting that 5-FU’s cytotoxic activity might be independent of its effects on deoxynucleotide pools (Figure 1E). Furthermore, when we focused on the apoptotic response, there was typically minimal benefit of the 5-FU-containing drug combination beyond the effects of the DNA-damaging drugs in isolation, and even examples of antagonism between the drugs where the addition of 5-FU decreased the cell death induced by oxaliplatin or irinotecan (Figure 1 G and Supplemental Figure 1).

To test whether this lack of synergistic killing observed in colon cancer cell lines was reflected in the progression-free survival of human colon cancer patients, we used publicly available data and calculated the expected additivity of progression-free survival (PFS) or time to disease progress (TTP) for patients treated with these drugs individually (Becouarn et al., 1998; Rougier et al., 1997) or in combination, and compared this to the observed PFS or TTP from publicly available FOLFOX or FOLFIRI trials (Becouarn et al., 1998; Douillard et al., 2000; Giacchetti et al., 2000; Rougier et al., 1997), (see Methods) (Hwangbo et al., 2022). Analysis of this clinical trial data revealed at best, an additive but non-synergistic effect for the combination of oxaliplatin with 5-FU (Figure 1H), and a sub-additive effect for the combination of 5-FU and irinotecan (Figure 1I). Taken together, these results suggest that the cytotoxic activities of 5-FU minimally enhanced the DNA damage-induced cytotoxicity conferred by these agents, and are entirely consistent with the beneficial effects of combination chemotherapy arising from patient-to-patient variability rather than drug additivity or synergy (Palmer and Sorger, 2017).

### 5-FU-dependent cytotoxicity in mammalian CRC cells is primarily dependent upon its incorporation into RNA

Although 5-FU’s primary mechanism of action has been classically ascribed to TYMS- and DDR-dependent activities, the results described above suggested that 5-FU’s DNA damage effects may not fully explain its efficacy. Indeed, we observed that TYMS mRNA expression across the CCLE cell line panel showed no correlation with 5-FU sensitivity (Supplemental Figure 2A), and its expression was unable to distinguish responders from non-responders in clinical trials of FOLFIRI or FOLFOX for colorectal cancer (Supplemental Figure 2B, C) (Del Rio et al., 2007; Tsuji et al., 2012). Furthermore, we found that maximal TYMS inhibition in our cell lines occurred at concentrations of 5- FU that do not induce apoptosis in short-term assays or affect proliferation over 72 hours (Supplemental Figure 2D-G). We therefore examined how CRC cells utilize 5-FU relative to uracil to better understand the global effects of 5-FU on nucleic acid metabolism. Uracil and 5-FU are metabolized by the same pyrimidine salvage pathway enzymes leading to their accumulation in both RNA and DNA (Longley et al., 2003) (Figure 2A). Recent reports have suggested that regulation of enzymes in this pathway is important, since perturbations that lead to greater 5-FUMP increase 5-FU’s potency (Cantor et al., 2017; Garcia-Gonzalez et al., 2017; Scott et al., 2017). Despite this, the ultimate fate of the drug remained unclear. Importantly, 5-FUMP is not committed to RNA incorporation, since the downstream metabolite, 5-FUDP, is metabolized to 5-FdUDP by ribonucleotide reductase, leading to its incorporation into DNA. We hypothesized that the relative importance of 5-FU’s nucleic acid effects would likely be a combination of the drug’s metabolic fate and the severity of the consequences of its incorporation into DNA, RNA, or both. Using ^14^C-labeled 5-FU, we found that far more of the drug accumulated in RNA than DNA, consistent with previous results (Pettersen et al., 2011), but this was both qualitatively and quantitatively distinct from how cells use ^14^C-labeled uracil (Figure 2B). ^14^C-uracil was distributed approximately equally between RNA and DNA (presumably as thymidine), but significantly more radioactivity was detected in RNA from ^14^C-5-FU-treated cells. We observed this difference in both complete and dialyzed serum, with no apparent difference in the initial uptake of the two compounds (Supplemental Figure 2H and I). This difference between 5-FU and uracil could potentially be explained by: 1) 5- FU-dependent effects on nucleotide metabolism and/or DNA replication altering how the cell uses uracil metabolites or 2) 5-FU and uracil having distinct metabolic/incorporation preferences due to inherent properties of the compounds – i.e. methyl versus fluoro- substituents in the 5 position of the pyrimidine ring. To distinguish between these possibilities, cells were treated with unlabeled 5-FU, and the subsequent nucleic acid incorporation of ^14^C-uracil was examined. As shown in Figure 2C, although 5-FU treatment decreased the amount of ^14^C-uracil incorporated into both RNA and DNA, it did not qualitatively alter the distribution of the labeled uracil, suggesting that 5-FU has an inherent priority over uracil for RNA incorporation. Given that the intracellular concentration of UTP is typically >500 μM (Traut, 1994), 5-FU’s ability to outcompete uracil is likely necessary for these RNA-dependent effects to be observed.

**Figure 2:**
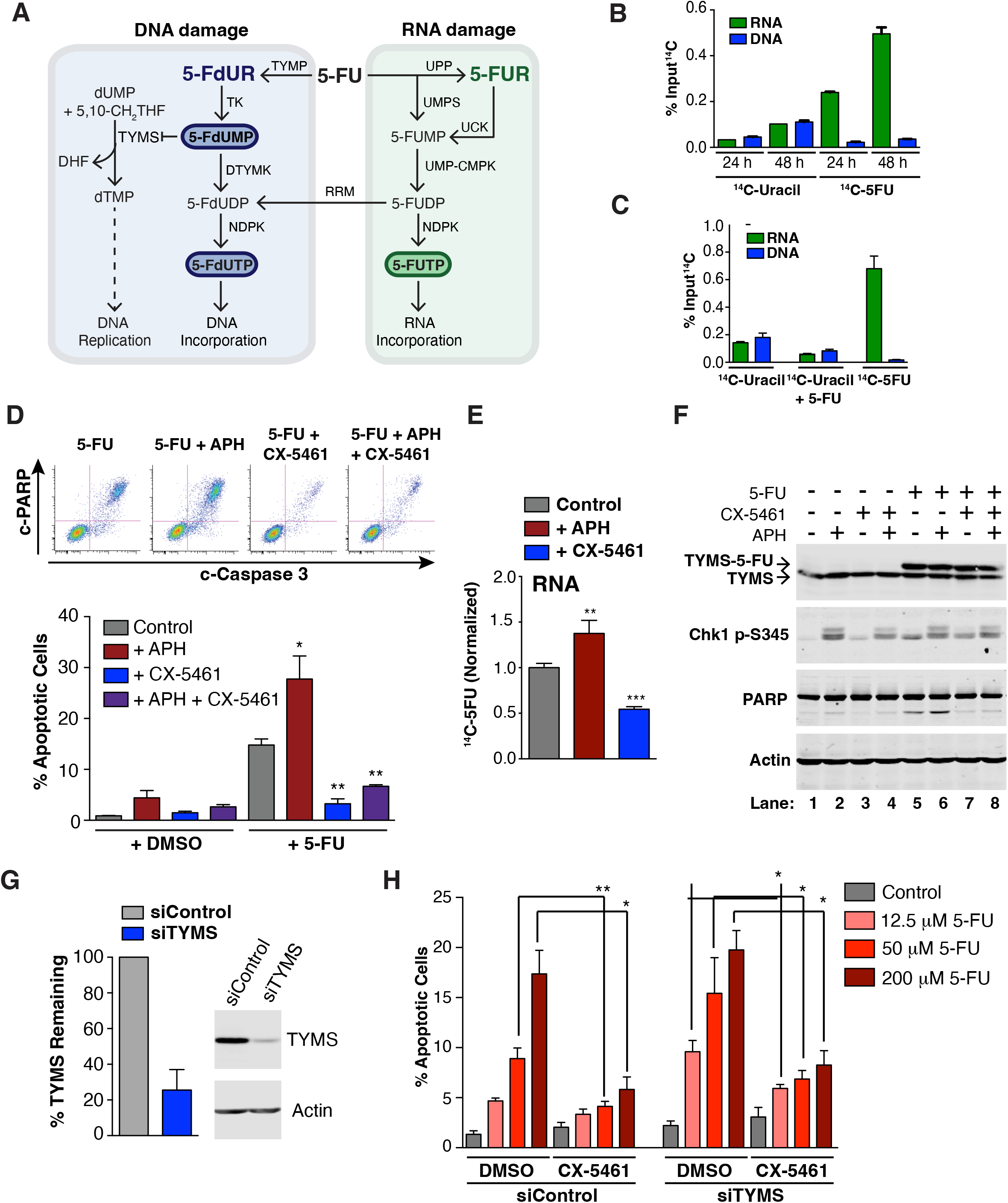
The canonical mechanism of 5-FU action does not explain drug-induced lethality. A) 5-FU is metabolized by the pyrimidine salvage pathway to generate multiple bioactive metabolites. B) HCT116 cells were treated with 0.6 μCi [2-^14^C]-5-FU or [2-^14^C]-uracil for 24 or 48 hours. RNA and DNA were purified and incorporation was quantified using a scintillation counter. C) HCT116 cells were treated with 0.6 μCi [2-^14^C]-5-FU, 0.6 μCi [2-^14^C]-uracil, or 0.6 μCi [2-^14^C]-uracil + 12.5 μM unlabeled 5-FU (the drug dose used when cells are treated with 0.6 μCi [2-^14^C]-5-FU). RNA and DNA were purified and incorporation was quantified as in panel B. D) HCT116 cells were treated with aphidicolin (APH) for 4 hours, CX-5461 for 1 hour, or both prior to addition of 200 μM 5-FU for 24 h. Cells were harvested, fixed, and the percentage of apoptotic cells was determined by flow cytometry. Representative FACS data were presented and quantified at bottom as mean ± s.e.m. from three independent experiments. \**P < 0.05,* \*\**P < 0.01*. E) HCT116 cells were treated with aphidicolin or CX-5461 as in D) prior to addition of 0.6 μCi [2-^14^C]-5-FU. RNA was purified and incorporation was quantified as in panel B. F) HCT116 cells were treated as in D), and collected after 24 h for western blot. G) HCT116 cells were transfected with siControl or siTYMS for 48 h, and knockdown efficiency was determined by western blot. H) HCT116 cells were transfected with siRNA as in G), and cells were treated with 5-FU for 24 hours. The percentage of apoptotic cells was quantified as mean ± s.e.m. from three independent experiments. \**P < 0.05,* \*\**P < 0.01*.

Although these experiments show that 5-FU has priority for RNA incorporation, it is unclear if RNA is the relevant target responsible for the drug’s lethality, or is merely a non-productive sink. To address this, we examined the effects of inhibiting RNA synthesis versus DNA synthesis on 5-FU-mediated apoptotic cell death. Given that rRNA represents the majority of cellular RNA and 5-FU’s previously described ability to impair rRNA biogenesis (Burger et al., 2010; Lum et al., 2004), we compared the effect of the RNA polymerase I inhibitor CX-5461 with that of aphidicolin, an inhibitor of the major replicative DNA polymerases. As shown in Figure 2D, pre-treatment with the RNA polymerase I inhibitor CX-5461 (Drygin et al., 2011) caused a dramatic *resistance* to 5- FU-induced apoptosis, while blocking DNA replication with aphidicolin unexpectedly *enhanced* 5-FU-induced cell death. These altered apoptotic effects upon 5-FU treatment directly correlated with its incorporation into RNA; CX-5461 pre-treatment resulted in a ∼50% decrease in 5-FU incorporation into RNA while aphidicolin pre-treatment increased the amount of 5-FU incorporated in RNA by ∼30% (Figure 2E). Furthermore, the enhanced apoptosis observed with aphidicolin pre-treatment was eliminated by blocking the enhanced 5- FU incorporation into RNA using co-pretreatment with both aphidicolin and CX-5461 (Figure 2D and F).

Interestingly, oxaliplatin treatment led to significantly reduced incorporation of ^14^C- 5-FU in RNA, suggesting that the sub-additivity that we observed in 5-FU/oxaliplatin combination might be due to blocking 5-FU’s RNA-dependent effects in a similar way as CX-5461 (Supplemental Figure 2J, K). This is consistent with the recent finding that oxaliplatin can act as a potent inhibitor of rRNA transcription, which significantly contributes to its mechanism of action (Bruno et al., 2017). Importantly, CX-5461 treatment did not suppress the ability of 5-FU to inhibit TYMS or activate the DNA damage response indicated as CHK1 phosphorylation at S345 residue (Figure 2F), indicating that RNA polymerase I inhibition does not block either the uptake of 5-FU or its DNA damage effects. Thus, treatments that increase the amount of 5-FU incorporation into RNA result in greater cytotoxicity whereas those that prevent the drug’s RNA incorporation provide resistance to cell death.

Because TYMS has long been considered a primary mechanistic target of 5-FU (Longley et al., 2003), it was particularly surprising that blocking DNA replication actually enhanced the drug’s apoptotic effect. It has previously been shown that TYMS knockdown sensitizes cells to 5-FU treatment (Muhale et al., 2011), consistent with TYMS as a direct 5-FU drug target. In agreement with this, we also observed a modest increase in cell death after 5-FU treatment upon TYMS knock-down, particularly at lower levels of 5-FU treatment where the total amount of death was small. However, at all 5-FU doses the extent of drug-induced cell death remained sensitive to CX-5461 treatment, even when TYMS was knocked down (Figure 2G and H). Thus, the effect of RNA polymerase inhibition was dominant even when the drug’s DNA damage effects were enhanced by TYMS knockdown. Therefore, although 5-FU is able to cause DNA damage, the drug’s efficacy in mammalian CRC cell killing appears to be primarily dependent upon its incorporation into RNA.

### 5-FU downstream metabolites biased towards RNA or DNA incorporation reveal that RNA damage more potently induces CRC cell death

To better dissect the phenotypic consequences of DNA- and RNA-dependent 5-FU effects, we used metabolites downstream of 5-FU: 5-fluorouridine (5-FUR) and 5-fluoro- 2’- deoxyuridine (5-FdUR)—that are strongly biased towards specific incorporation into RNA or DNA, respectively (Figure 2A) (Pettersen et al., 2011). In agreement with these distinct metabolic fates, disruption of RNA integrity by 5-FUR treatment did not activate the DNA damage response, as assessed by time-dependent phosphorylation of the DNA damage checkpoint effector kinases Chk1 and Chk2, whereas both 5-FU and 5-FdUR treatment resulted in robust activation (Figure 3A). However, despite its inability to activate the DDR, 5-FUR was a far more potent inducer of apoptosis compared to 5-FdUR in both a p53 WT and an isogenic p53-null cell line (Figure 3B and C). Furthermore, despite both causing equivalent stabilization of p53, 5-FUR treatment resulted in a far more robust accumulation of PUMA than 5-FdUR. (Figure 3D)

**Figure 3:**
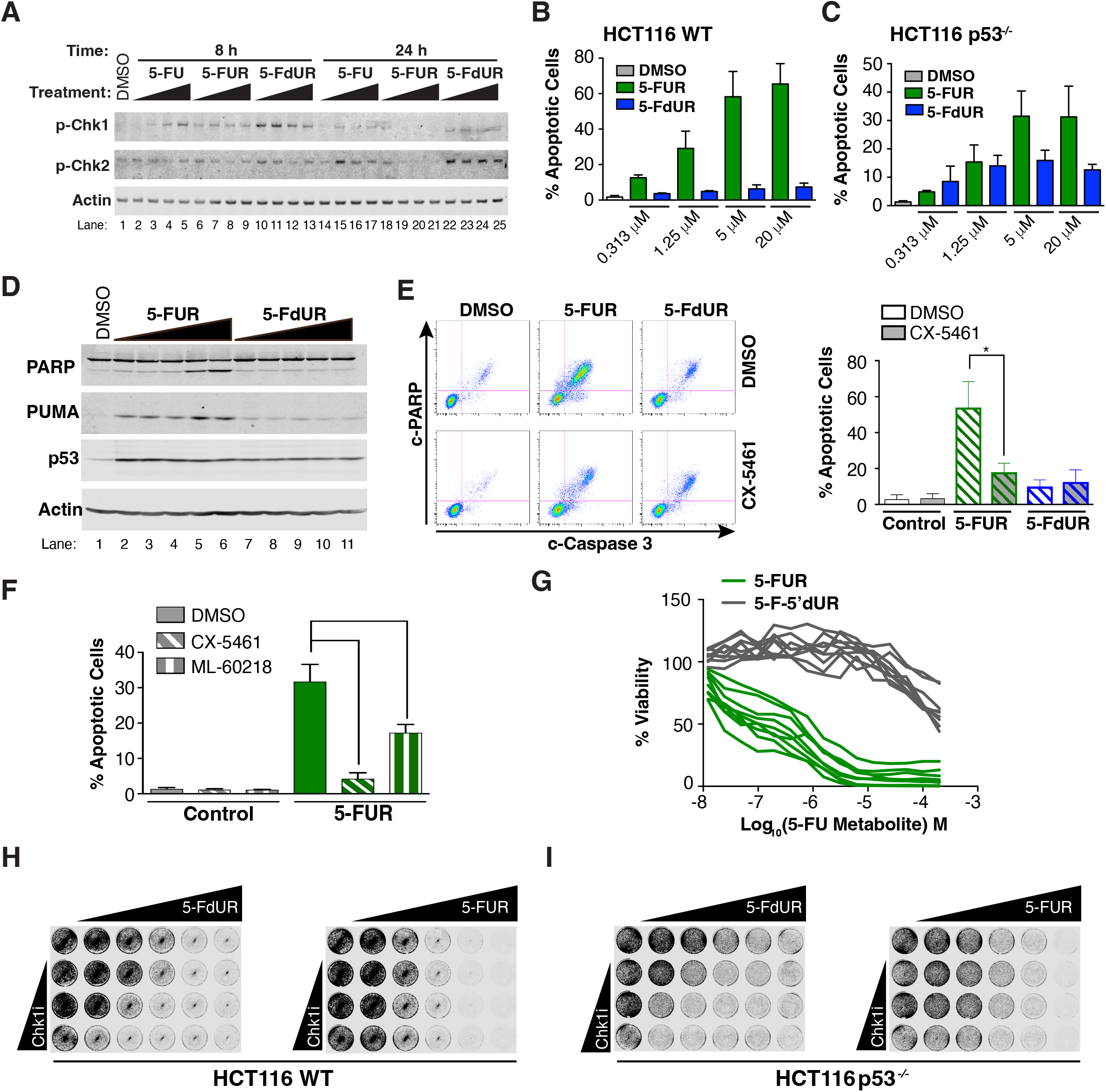
5-FU metabolites biased towards RNA and DNA damage reveal pathway specific responses. A) HCT116 cells were treated with equal concentrations of 5-FU, 5-FdUR, or 5-FUR for 8 or 24 hours, and activation of the DDR was monitored by detecting phosphorylation of Chk1 and Chk2. B) Response to 5-FUR and 5-FdUR in HCT116 cells was determined by flow cytometry 24 hours after drug treatment C) As in B), except with HCT116 *p53^-/-^* cells after 48 hours of treatment. D) SW48 cells were treated with increasing doses of 5-FUR or 5-FdUR (10 μM top concentration) for 16 hours, and lysates were analyzed by western blot. E) HCT116 cells were treated with CX-5461 prior to addition of 2 μM 5-FUR or 20 μM 5- FdUR. Induction of apoptosis was determined by flow cytometry 24 hours after drug treatment. Representative FACS data were presented and quantified at right as mean ± s.e.m. from three independent experiments. \**P < 0.05*. F) HCT116 cells were treated with CX-5461 or the RNA Polymerase III inhibitor ML-60218 prior to 5-FUR addition. After 4 hours of 5-FUR treatment, drugs were washed out to reduce the toxicity of ML-60218 and viability was determined by flow cytometry. G) A CRC cell line panel (see Figure 4A) was treated with 5-FUR or 5-F-5’dUR for 72 hours, and viability was determined by CellTiter-Glo. Each line represents the dose- response curve for 5-FUR or 5-F-5’dUR. H) HCT116 WT cells were treated with a dose-response matrix of a Chk1 inhibitor and 5- FdUR (left) or 5-FUR (right). Viability following treatment was determined by staining with SYTO60 I) As in H), except using HCT116 *p53^-/-^* cells.

As observed previously with 5-FU (Figure 2D), cell cytotoxicity following 5-FUR treatment was dependent upon incorporation of the metabolite into RNA. Cell death was found to be dramatically reduced if the cells were pre-treated with CX-5461 prior to 5- FUR administration (Figure 3E). In marked contrast, RNA polymerase I inhibition did not block the modest amount of apoptosis caused by 5-FdUR treatment. Interestingly, RNA polymerase III inhibition using ML-60218 (Wu et al., 2003) was also capable of reducing the amount of 5-FUR-induced apoptosis (Figure 3F), albeit to a lesser extent than CX- 5461. To further investigate a requirement for RNA incorporation in the induction of cell death, we examined whether replacing the 5’-OH of 5-FUR with a methyl group (5-F- 5’dUR), which blocks its utilization by RNA polymerases, affected 5-FUR’s cytotoxic activity. As shown in Figure 3G, 5-F-5’dUR had little effect on cell viability relative to 5- FUR, further demonstrating a dependency on RNA incorporation. These results, therefore, indicate that 5-FU’s RNA damage effects are an intrinsically more potent inducer of cell death compared to its DDR-dependent effects in CRC cells.

Most of the chemical or genetic perturbations reported to date that increase tumor cell sensitivity to 5-FU have focused on enhancing the drug’s effects on DNA damage (Muhale et al., 2011; Xiao et al., 2013). To independently examine this, the effects of Chk1 inhibition or TYMS down-regulation on the response to 5-FUR versus 5-FdUR treatment were investigated. As shown in Figures 3H and I, inhibition of Chk1 significantly enhanced the response to 5-FdUR in both p53 wild-type and null cells while having little effect on the more potent response of these cells to 5-FUR. Likewise, TYMS knockdown more strongly enhanced cell sensitivity towards 5-FdUR than to 5-FUR (Supplemental Figure 3), further suggesting that 5-FUR and 5-FdUR are biased towards mediating the drug’s RNA and DNA damage effects, respectively, and that known 5-FU synergistic treatments have primarily targeted the DNA damage-dependent mechanism of action of the drug.

### 5-FUR specifically triggers ubiquitination of multiple ribosomal proteins

The results shown above suggest that the 5-FU metabolites, 5-FdUR and 5-FUR have distinct phenotypic effects on tumor cells mediated through DNA or RNA damage, respectively. Unlike the well-studied DNA damage response, relatively little is known about the signaling events downstream of RNA damage. To explore this, we performed a comprehensive mass-spectrometry-based analysis of the phospho-SQ/TQ proteome, total phospho-proteome and ubiquitin-proteome using quantitative 11-plex TMT mass spectrometry in CRC cells treated with 5-FdUR or 5-FUR (Methods, Figure 4A). To detect the most direct effects of each compound, we chose an acute time point (6 hours) following accumulation into RNA and DNA respectively, that correlated with increased p53 accumulation in 5-FUR treated cells and KAP1 phosphorylation in 5-FdUR treated cells by western blotting (Figure 4B). In our analysis, we sought changes in post-translational modifications that were unique or preferentially enriched in one treatment compared to the other since we hypothesized such changes would be most likely to explain the differences in phenotypic effects of the two 5-FU metabolites.

**Figure 4:**
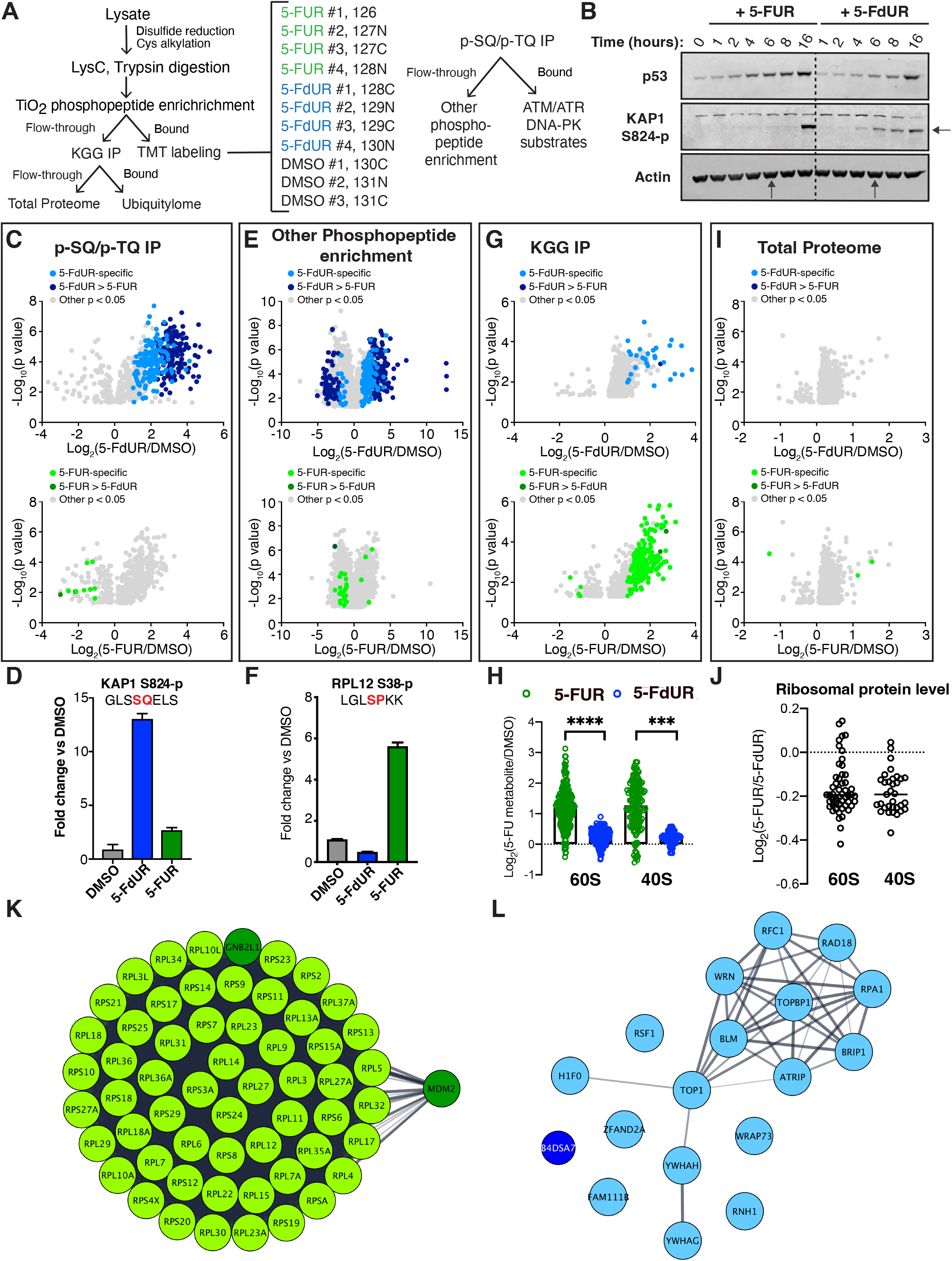
Distinct signaling responses exist to 5-FU metabolites that cause DNA and RNA damage. A) Schematic for 11-plex TMT mass spectrometry to assess signaling response to 5-FUR and 5-FdUR. B) HCT116 cells were treated with 1 μM 5-FUR or 5-FdUR for the indicated times, and the indicated proteins were assessed by western blot. C) Volcano plot for p-SQ/p-TQ IP showing all peptides that were significantly different in 5-FdUR (top) or 5-FUR (bottom) treated cells compared to cells treated with DMSO. Peptides were classified as being specific or enriched for one metabolite as described in materials and methods. D) Quantification of KAP1 S824-p peptide from p-SQ/p-TQ IP across all treatment conditions as an example of a 5-FdUR phosphopeptide. E) As in C), showing significant peptides identified following phosphopeptide enrichment. F) Quantification of RPL12 S38-p peptide from all treatment conditions as an example of a 5-FUR specific phosphopeptide. G) As in C), showing significant peptides identified following KGG IP. H) Quantification of peptides identified following KGG IP from proteins that are part of the 60S or 40S ribosomal subunits for 5-FUR or 5-FdUR treatment. I) As in C), showing proteins whose expression significantly changed following compound treatment. J) Total levels of ribosomal proteins decrease following treatment with 5-FUR relative to 5-FdUR. K) STRING functional protein association networks of 5-FUR-induced ubiquitinated proteins generated by Cytoscape software. L) STRING functional protein association networks of 5-FdUR-induced ubiquitinated proteins generated by Cytoscape software.

The DNA damage response kinases ATM, ATR, and DNA-PK preferentially phosphorylate SQ/TQ sites, and p-SQ/p-TQ motif antibodies have been used previously to enrich for substrates of these kinases after ionizing radiation and other genotoxic treatments (Kim et al., 1999; Matsuoka et al., 2007; O’Neill et al., 2000). As shown in Figure 4C and D, 5-FdUR treatment resulted in a marked upregulation of SQ/TQ phophopeptide abundance, whereas 5-FUR had a far more muted effect (Figure 4C and D). Likewise, while both treatments caused changes more broadly to the total phospho-proteome, relatively few changes were specific to the 5-FUR treated cells compared to 5- FdUR (Figure 4E and F). Interestingly, among the few phosphoproteins whose levels were uniquely changed by 5-FUR treatment are a ribosomal protein (RPL12) and an RNA pseudouridine synthase (RPUSD2) implicated in ribosome biogenesis. In contrast, a large number of the phosphorylated proteins containing 5-FdUR-modulated sites have been previously implicated in DNA repair and replication, consistent with strong activation of the DNA damage response and replication stress by this compound (Supplemental Figure 4).

Strikingly, KGG enrichment identified many ubiquitinated peptides following 5-FUR treatment and far fewer following 5-FdUR treatment (Figure 4G). Nearly all of the unique, significantly enhanced ubiquitinated peptides following 5-FUR treatment were identified as belonging to 40S and 60S ribosomal proteins (Figure 4H and K). This effect was specific to 5-FUR, with 5-FdUR treatment generating many fewer differentially abundant ubiquitination events, most of which occurred on proteins involved in the DNA damage response (Figure 4L). Furthermore, nearly all of the ribosomal proteins that were detected by mass spectrometry had at least one lysine that was differentially ubiquitinated following 5-FUR treatment, suggesting that the ribosome is a key signaling hub for the effects of 5- FUR. While the overall change in the total proteome was relatively small at this time point, there was a strongly significant relative decrease in the levels of ribosomal proteins in cells treated with 5-FUR compared to 5-FdUR (Figure 4I and J).

Overall, this data indicates that, while there are some common alterations in the total and modified proteome induced by treatment with these two 5-FU metabolites – potentially reflecting some level of interconversion of RNA and DNA damaging metabolites – each compound has unique consequences on cellular signaling, likely leading to differences in phenotypic effects.

### 5-FUR causes proteasome-dependent, but not lysosome-dependent, degradation of ribosomal proteins

To examine whether the decrease in ribosomal protein levels following 5-FUR treatment reflects loss or redistribution of intact ribosomes compared to 5-FdUR, we used sucrose gradient centrifugation to profile ribosomes in 5-FUR and 5-FdUR-treated cells. As shown in Figure 5A, 5-FUR treatment eliminated both the polysome and monosome fraction as well as depleting the levels of the individual 40S and 60S subunits. In contrast, 5-FdUR treatment primarily shifted the translation-proficient polysomes to monosomes and free ribosomal subunits with no decrease in the total level of ribosomal RNA. These findings indicate that 5-FUR treatment results in ribosome degradation, while 5-FdUR interferes with the translation process but does not alter total ribosome abundance. Furthermore, this 5-FUR-induced loss was specific to ribosomes, since no changes were observed in the levels of markers for other subcellular organelles (Figure 5B).

**Figure 5:**
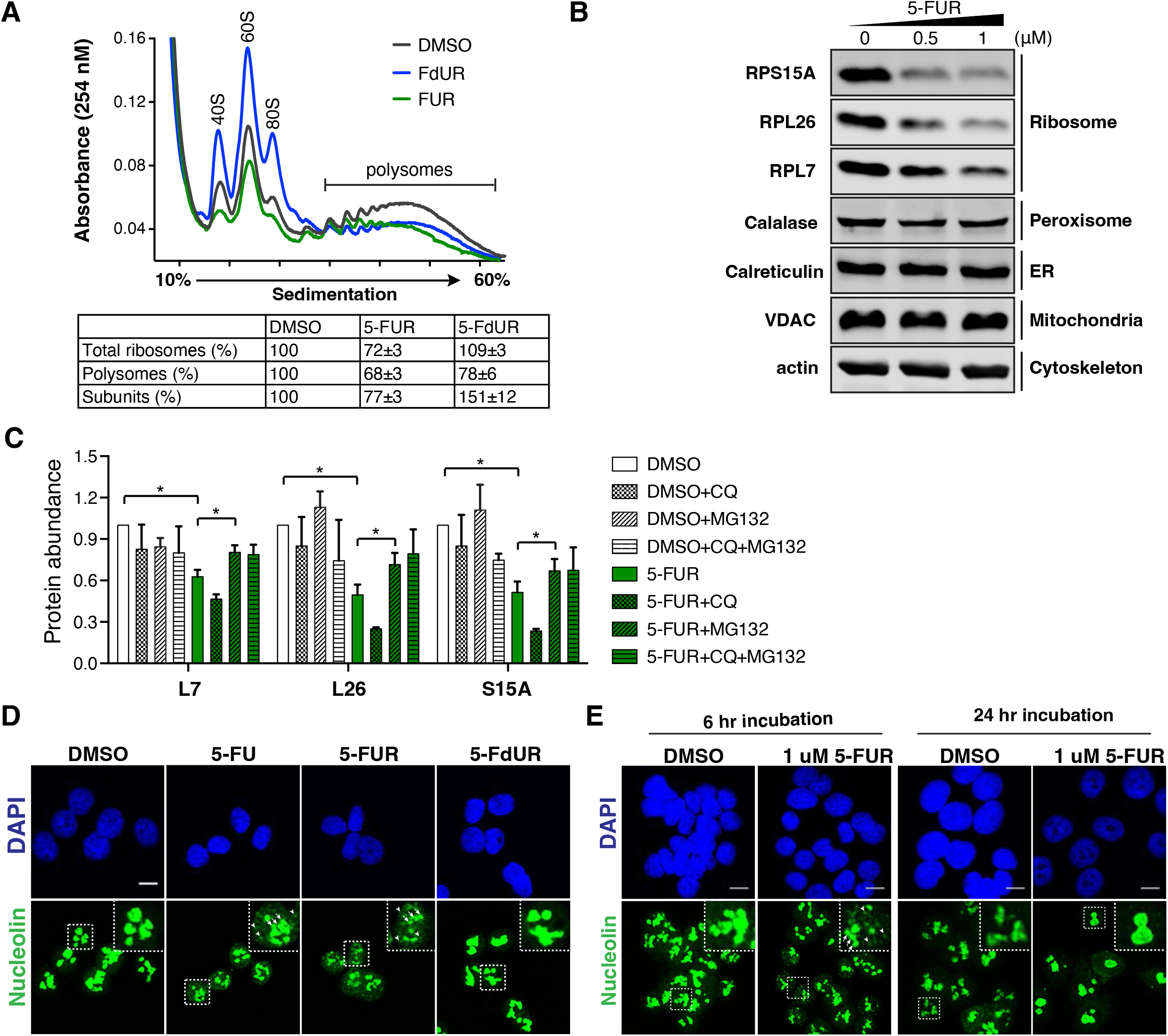
5-FUR causes proteasome-dependent degradation of ribosomal proteins. A) HCT116 cells were treated with 2 μM 5-FUR or 2 μM 5-FdUR for 18 hours and subjected to polysome gradient to reveal ribosome profile. B) HCT116 cells were treated with increasing concentration of 5-FUR for 18 hours. The total cell lysates were prepared using Laemmli buffer and subjected to immunoblotting against the indicated antibodies. C) HCT116 cells were treated with 2 μM 5-FUR together with 20 uM chloroquine or 5 uM MG132 for 18 hours. Data from three independent experiments were quantified as mean ± s.e.m. \**P < 0.05*. D) HCT116 cells were treated with DMSO, 80 μM 5-FU, 2 μM 5-FUR, or 20 μM 5-FUR for 6 hours and subjected to immunofluorescence against nucleolin. The arrow indicates necklace-like nucleolin distribution, and the arrowhead indicates punctate nucleolin foci. The scale bar is 10 μm. E) HCT116 cells were treated with DMSSO or 1 μM 5-FUR for 6 and 24 hours and subjected to immunofluorescence as shown in 5D.

In response to various stimuli including starvation, oxidative stress, and inhibition of translation with the mTOR inhibitors rapamycin and Torin-1, ribosomes are known to undergo a specific autophagic process, ribophagy, that involves ubiquitination- and lysosomal-dependent degradation (Kocaturk and Gozuacik, 2018). To examine whether the ubiquitination of ribosomal proteins and loss of total ribosomes induced by 5-FUR reflected ribophagy, HCT116 cells were treated with 5-FUR and co-incubated with either chloroquine, which inhibits autophagy by blocking the fusion of lysosomes and autophagosomes (Mauthe et al., 2018), or with the proteasome inhibitor MG132. Surprisingly, chloroquine co-treatment resulted in a further decrease in ribosome abundance compared to 5-FUR treatment alone (Supplemental Figure 5A), as well as a decrease in the levels of ribosomal proteins (Figure 5C and Supplemental Figure 5B). In contrast, treatment with the proteasome inhibitor MG132 partially rescued the protein levels of these ribosomal proteins (Figure 5C), indicating that 5-FUR-induced ubiquitination of ribosomal proteins results in their degradation through a proteasome-dependent pathway rather than through lysosome-dependent ribophagy.

This autophagosome-independent degradation of ribosomal proteins suggests that in colorectal cancer cells 5-FU and 5-FUR likely exert their primary effect during the process of ribosome biogenesis rather than causing stress-dependent autophagy of pre-existing ribosome pools, consistent with previous findings related to 5-FU-induced cell death in yeast (Lum et al., 2004), and in mammalian cells (Ghoshal and Jacob, 1994; Kanamaru et al., 1986). To further examine this, we stained cells for nucleolin, an abundant protein involved in ribosome biogenesis that serves as a marker of nucleolar morphology (Ginisty et al., 1999). As shown in Figures 5D and E, treatment with 5-FU and 5-FUR, but not with 5-FdUR, disrupted the normal nucleolar architecture resulting in the appearance of small punctate nucleolin foci and nucleolar necklaces within 6 hours of treatment, and large nucleolar aggregates by 24 hours, consistent with nucleolar restructuring and defect in ribosome biogenesis. (Latonen, 2019)

### 5-FUR induces lysosomal-dependent autophagy of ribosomal rRNA

We next examined the effect of 5-FU, 5-FUR and 5-FdUR on the total levels of ribosomal RNA within the cells. Consistent with the impairment of ribosome biogenesis, 5-FU and 5-FUR, but not 5-FdUR treatment caused a 20-30% decrease in the amount of 18S and 28S rRNA in whole cell extracts when analysed by agarose gel electrophoresis. (Figure 6A). In contrast, 5-FdUR treatment actually resulted in a slightly increase in 18S and 28S rRNA abundance likely as a consequence of inhibiting cell division leading to an increase in cell size (Supplemental Figure 6A). Similarly, 5-FUR, but not 5-FdUR treatment caused a significant decrease in the amount of 18S, 28S and 45S rRNA when analysed by quantitative PCR (Figure 6B).

**Figure 6.**
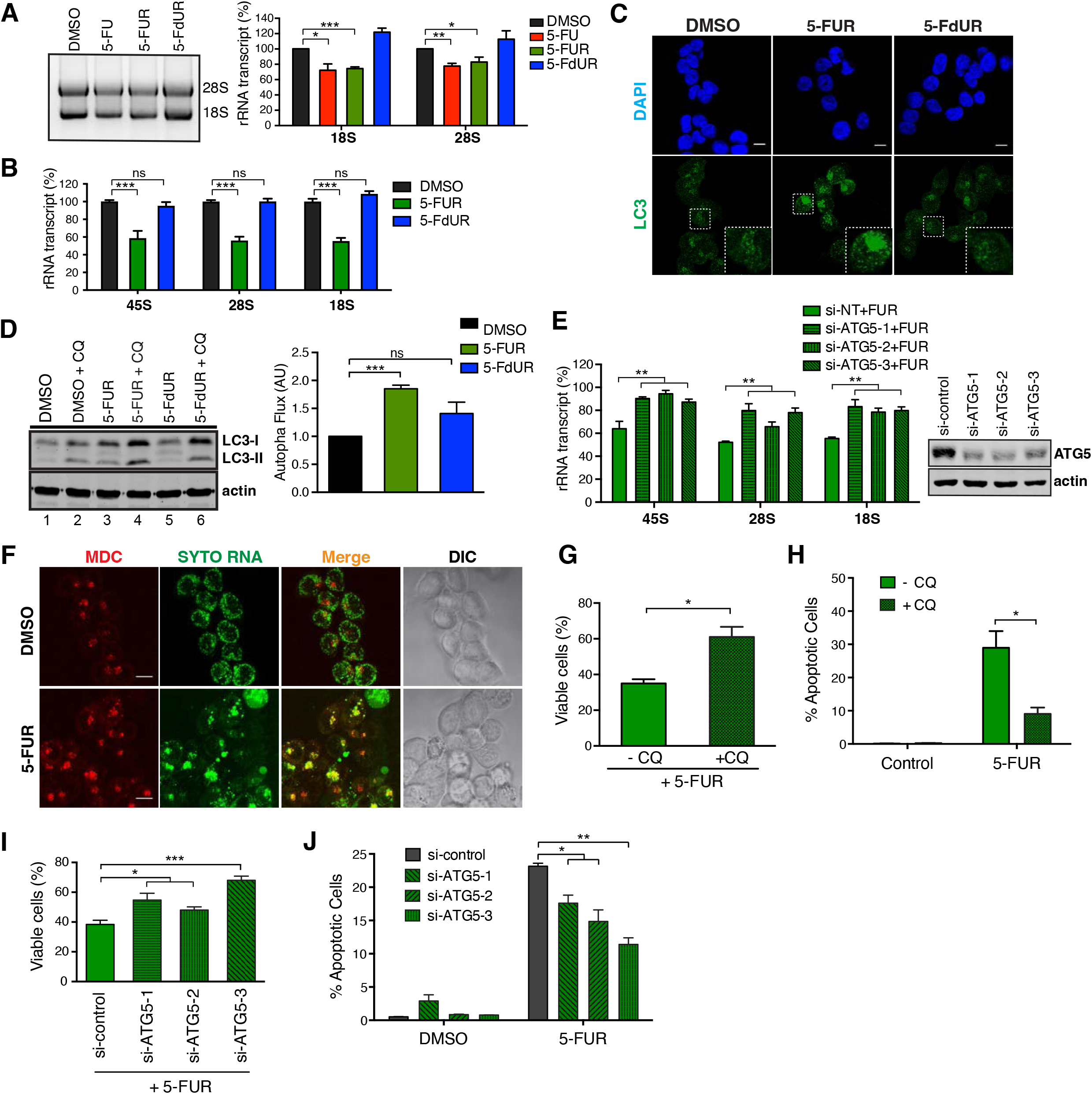
5-FUR causes autophagy-dependent degradation of ribosomal rRNA. A) HCT116 cells were treated with 40 μM 5-FU, 2 μM 5-FUR or 2 μM 5-FdUR for 18 hours. The extracted RNA was subjected to electrophoresis to show the 18S/28S ribosomal RNA abundance. 18S and 28S rRNA abundance were quantified at right as mean ± s.e.m. from three independent experiments. \**P < 0.05,* \*\**P < 0.01,* \*\*\**P < 0.001*. B) HCT116 cells were treated with 2 μM 5-FUR or 2 μM 5-FdUR for 18 hours. The extracted RNA was reverse-transcribed to cDNA using random hexamer and subjected to qPCR analysis to show the 18S, 28S and 45S ribosomal RNA abundance. Data is presented as mean ± s.e.m. from three independent experiments \*\*\**P < 0.001,* ns, not significant (*P > 0.05*). C) HCT116 cells were treated with 2 μM 5-FUR or 2 μM 5-FdUR for 18 hours and subjected to immunofluorescence against indicated antibodies. Representative fluorescent images of LC3 expression of three independent experiments. The scale bar is 10 μm. D) HCT116 cells were treated with 2 μM 5-FUR or 2 μM 5-FdUR for 15 hours. 20uM chloroquine (CQ) was added for another 3 hours. LC3-II was normalized to actin and autophagic flux was calculated by subtracting the value of normalized LC3-II in the presence of CQ by that without CQ. Autophagy flux of DMSO-treated cells was set to 1, and the rest of the samples were normalized accordingly. Data is presented as mean ± s.e.m. from three independent experiments. \*\*\**P < 0.001*. E) HCT116 cells were transfected with non-targeting or ATG5 siRNA and then treated with or without 2 μM 5-FUR for 18 hours. The extracted RNA was reverse-transcribed to cDNA and subjected to quantitative PCR against individual ribosomal RNA transcript. Data was normalized to DMSO-treated cells and presented as mean ± s.e.m. from three independent experiments. \*\**P < 0.01*. F) HCT116 cells were treated with DMSO or 2 μM 5-FUR for 12 hours. After the incubation, live cells were stained with SYTO RNA select for 20 min at 37 °C, and then with 50 μM monodansylcadaverine (MDC) for a further 30 min at 37 °C. The data showed representative fluorescent images of MDC and SYTO RNA staining of three independent experiments. DIC, differential interference contrast. The scale bar is 10 μm. G) HCT116 cells were treated with 2 μM 5-FUR in the presence or absence of 20 μM chloroquine (CQ) for 18 hours. Viability was accessed by trypan exclusion assay and each bar represents ratio of cells treated with 5-FUR to cells treated with DMSO in the presence or absence of chloroquine. Data was presented as mean ± s.e.m. from three independent experiments. \**P < 0.05*. H) HCT116 cells were treated with 2 μM 5-FUR in the presence of 20 μM chloroquine (CQ) for 24 hours. Induction of apoptosis was determined by flow cytometry and presented as mean ± s.e.m. from three independent experiments. \**P < 0.05*. I) HCT116 cells were transfected with non-targeting or ATG5 siRNA and then treated with or without 2 μM 5-FUR for 18 hours. Cell viability was determined as in Figure 6F. J) Non-targeting or ATG5 siRNA-transfected cells were treated with or without 1 μM 5- FUR for 20 hours. Induction of apoptosis was determined as in Figure 6G.

Autophagy is a well-recognized mechanism for degradation of damaged molecules in addition to intact organelles and macromolecular structures (Leidal et al., 2018). Although autophagy does not appear to be involved in the degradation of ribosomal proteins after 5-FU or 5-FUR treatment (Fig 5C), we wondered whether it played a role in the degradation of damaged ribosomal RNA. During the autophagy process, microtubule-associated protein 1A/1B-light chain 3 (LC3) is converted to a phosphatidylethanolamine (PE) membrane-localized form (LC3-II) to initiate the formation and lengthening of the autophagosome, which later fuses with the lysosome to degrade cargo inside the autophagosome (Gatica et al., 2018). To explore whether autophagy was involved in the degradation of rRNA following 5-FU and 5-FUR treatment, the localization and post-translational modification of LC3 was examined by immunofluorescence or immunoblotting, respectively. As shown in Figure 6C and D, 5-FUR treatment, but not 5- FdUR, resulted in the appearance of bright LC3-positive puncta and clusters and an LC3- II immuno-reactive band (Figure 6D, compare lane 3 with lanes 1 and 5). To further implicate autophagy in the cellular response to 5-FUR treatment, the flux through the autophagy pathway was quantified by measuring the increase in LC3-II accumulation 3 hours following the addition of chloroquine (Figure 6D), which blocks the fusion of lysosomes to the autophagosome (Chittaranjan et al., 2015). As quantified in the right panel, autophagic flux was statistically significantly increased by treatment with 5-FUR, but not by 5-FdUR.

To further examine the role of autophagy in rRNA degradation, RNA interference was used to target ATG5, a key protein necessary for conjugation of LC3-I with PE to form membrane-associated LC3-II (Otomo et al., 2013). Using 3 distinct siRNAs, ATG5 knockdown was observed to blunt the decline in 45S, 28S and 18S ribosomal RNAs following treatment with 5-FUR (Figure 6E). Likewise, chloroquine treatment also decreases the 5-FUR-induced decline of 45S, 28S and 18S ribosomal RNAs. (Supplemental Figure 6B). In addition, 5-FUR caused the accumulation of RNA-containing bodies that localized within autophagosomes as revealed by monodansylcadaverine (MDC) staining (Figure 6F). Taken together, these data indicated that 5-FUR treatment targets ribosomal RNA to autophagosomes by increasing autophagic flux, while targeting ribosomal proteins to the proteasome for degradation.

To explore the contribution of rRNA degradation to 5-FUR-induced tumor cell death, cells were treated with chloroquine to inhibit autophagosome-lysosome fusion. In the presence of 5-FUR, chloroquine treatment resulted in ∼2-fold increase in cell survival (Figure 6G and Supplemental Figure 6C). A focused examination of apoptotic cell death by co-staining for cPARP1 and cCaspase-3 revealed a similar ∼3-fold decrease in 5-FUR- induced apoptosis following chloroquine treatment (Figure 6H). Similarly, siRNA knockdown of ATG-5 also resulted in enhanced cell viability and reduced apoptotic cell death following 5-FUR treatment (Figure. 6I, J and Supplemental Figure 6D). Thus, inhibition of autophagic ribosomal RNA degradation following 5-FUR treatment reduces apoptotic cell death.

### KDM2A depletion enhances ribosomal RNA damage and synergizes with 5-FU to promote CRC tumor cell death

Since 5-FU induced cell death occurred primarily through fluorouridine incorporation into rRNA, inducing an RNA damage response with distinct modes of RNA and ribosomal protein degradation, we hypothesized that co-treatments inducing up-regulation of rRNA transcription might provide a potential approach to enhance the toxicity of 5-FU in cancer cells. Cells were therefore examined for their response to 5-FUR treatment when cultured in the presence or absence of glucose, which is a well-known regulator of rRNA transcription (Mariappan et al., 2011). Indeed, when cells were cultured in DMEM containing glucose, which enhanced nascent RNA transcription, they displayed significantly increased cytotoxicity to 5-FUR (Figure 7A, B). In addition, inhibition of mTORC1 that is known to positively regulate multiple steps in ribosome biogenesis (Iadevaia et al., 2014) greatly decreased the cytotoxicity to 5-FUR (Supplemental Figure 7A).

**Figure 7.**
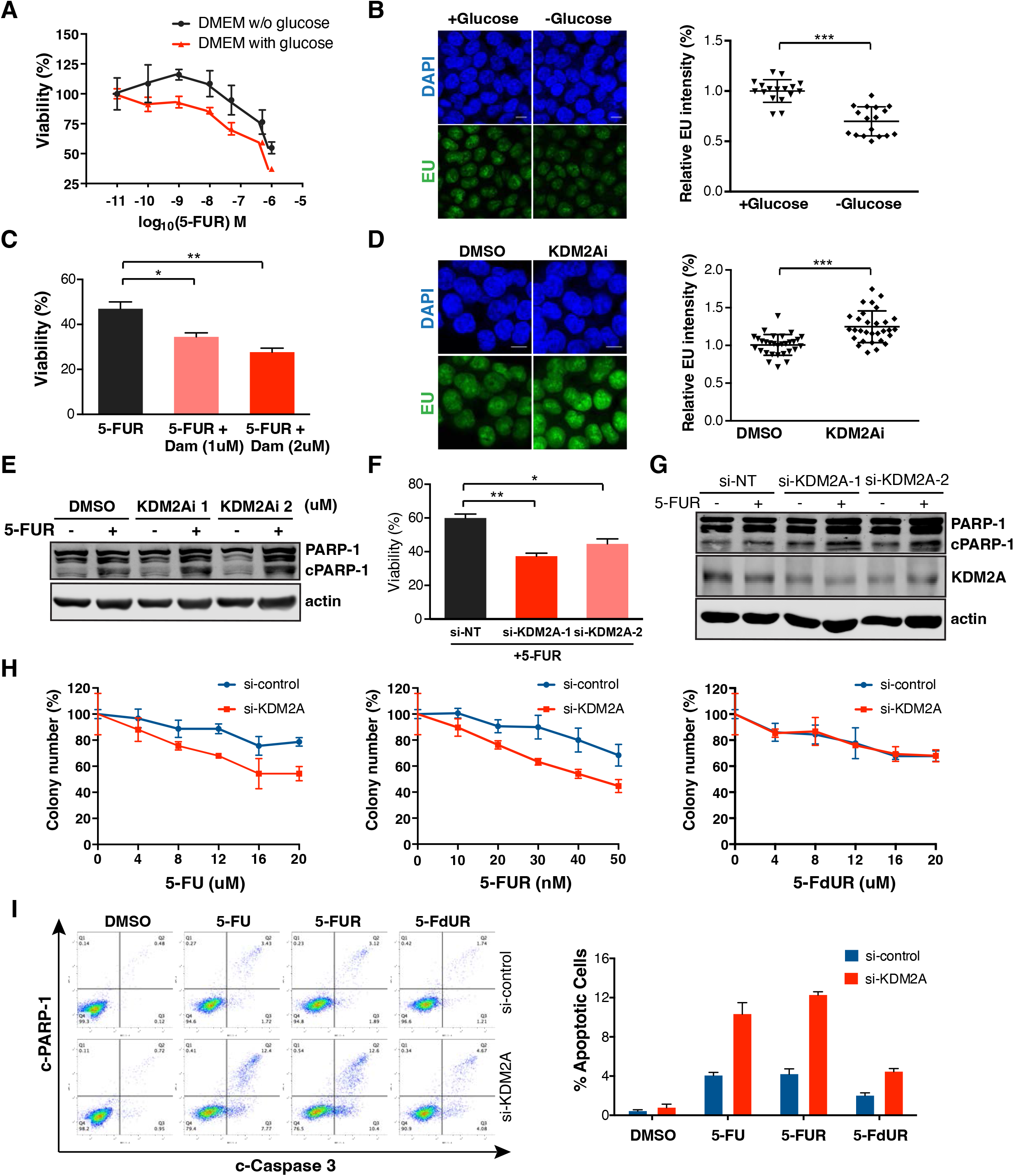
KDM2A depletion enhances RNA damage and synergizes cells to 5-FU. A) HCT116 cells were cultured in DMEM with or without glucose and treated with 5-FUR. 48 hours after 5-FUR treatment, viability was assessed by MTT assay. (representative of two independent experiments). B) HCT116 cells were cultured in DMEM with or without glucose for 24 hours followed by EU labelling and the newly synthesized RNA was analysed by immunofluorescence. The fluorescence intensity was quantified at right, each dot represents the average nuclear EU intensity for cells in a field. Data is presented as mean ± s.e.m. from two independent experiments. \*\*\**P < 0.001.* The scale bar is 10 μm. C) HCT116 cells were treated with 0.4 μM of 5-FUR in the presence or absence of Daminozide (KDM2A inhibitor). Viability was accessed by trypan exclusion assay and each bar represents ratio of cells treated with 5-FUR to cells treated with DMSO in the presence or absence of Daminozide. Data is presented as mean ± s.e.m. from three independent experiments. \**P < 0.05,* \*\**P < 0.01.* The scale bar is 10 μm. D) HCT116 cells were treated with DMSO or 2 μM Daminozide for 24 hours and following EU labelling as B). Data is presented as mean ± s.e.m. from three independent experiments. \*\*\**P < 0.001*. E) HCT116 cells were treated DMSO or 5-FUR in the presence or absence of Daminozide for 24 hours. The total cell lysates were prepared using Laemmli buffer and subjected to immunoblotting against the antibodies indicated. F) HCT 116 cells were transfected with non-targeting or KDM2A siRNAs and then treated with or without 0.4 μM 5-FUR for 24 hours. Viability was measured as shown in C). Data is presented as mean ± s.e.m. from three independent experiments. \**P < 0.05,* \*\**P < 0.01*. G) HCT 116 cells were transfected with non-targeting or KDM2A siRNA and then treated with or without 0.4 μM 5-FUR for 24 hours. The total cell lysates were prepared using Laemmli buffer and subjected to immunoblotting against the antibodies indicated. H) HCT 116 cells were transfected with non-targeting or KDM2A siRNAs and then treated with or without various concentration of 5-FU, 5-FUR and 5-FdUR for 24 hours. Viability was accessed by colony formation assay. (representative of three independent experiments). I). HCT 116 cells were transfected with non-targeting or KDM2A siRNAs and then treated with or without 40 μM 5-FU, 0.4 μM 5-FUR and 20 μM 5-FdUR for 24 hours. Induction of apoptosis was determined by flow cytometry. Representative FACS data were presented and quantified at right as mean ± s.e.m. from three independent experiments.

KDM2A is a recently described Jumonji domain-containing lysine demethylase that negatively regulates rRNA transcription by demethylating H3K36 on the rDNA promoter (Liu et al., 2021; Okamoto et al., 2019; Tanaka et al., 2010; Tanaka et al., 2015). Treatment of cells with daminozide, a selective KDM2A small molecule inhibitor (Rose et al., 2012), resulted in enhanced nascent RNA transcription and increased both 5-FUR toxicity and apoptotic cell death, as indicated by PARP-1 cleavage (Figure 7C-E). Furthermore, depletion of KDM2A using si-RNA resulted in a similar enhancement of 5- FUR-induced cytotoxicity and apoptosis (Figure 7F, G). To further test whether KDM2A depletion synergistically enhanced cell death in response to 5-FU-induced RNA damage, but not DNA damage, we compared the response of control and KDM2A-depleted HCT116 cells to 5-FU, 5-FUR (the 5-FU RNA damaging metabolite), or 5-FdUR (the 5- FU DNA damaging metabolite). As shown in Figures 7H and I, KDM2A depletion specifically sensitized HCT116 cells to 5-FU and 5-FUR, but not to 5-FdUR treatment. These data further implicate interference with ribosome biogenesis as a major mechanism for 5-FU-mediated cytotoxicity and suggest that therapeutic targeting of rRNA transcription pathways can be used to enhance 5-FU therapeutic responses.

### Sensitivity to RNA and DNA damage metabolites varies in a tissue-specific manner

In the isogenic HCT-116 cell lines tested above, 5-FUR was a significantly more potent inducer of apoptosis than 5-FdUR (Figure 3B and C). To more broadly examine the relative cytotoxic effects of these RNA and DNA damaging metabolites across a panel of CRC cell lines, we measured the GI_50_ (dose required for 50% inhibition of proliferation) for each compound and calculated the log_2_(GI_50_ 5-FUR/GI_50_ 5-FdUR) ratio as a metric of relative sensitivity for each cell line. This allowed us to examine the consequences of RNA and DNA damage independent of a cell line’s general sensitivity to cytotoxic agents. A ratio of zero indicates equal sensitivity towards the two compounds whereas a negative or positive number indicates greater sensitivity for 5-FUR or 5-FdUR, respectively. In every CRC cell line tested, we found that 5-FUR was a more potent inhibitor of cell viability than 5-FdUR (Figure 8A), indicating that CRC cells are broadly more sensitive to 5-FU’s RNA damaging effects.

**Figure 8:**
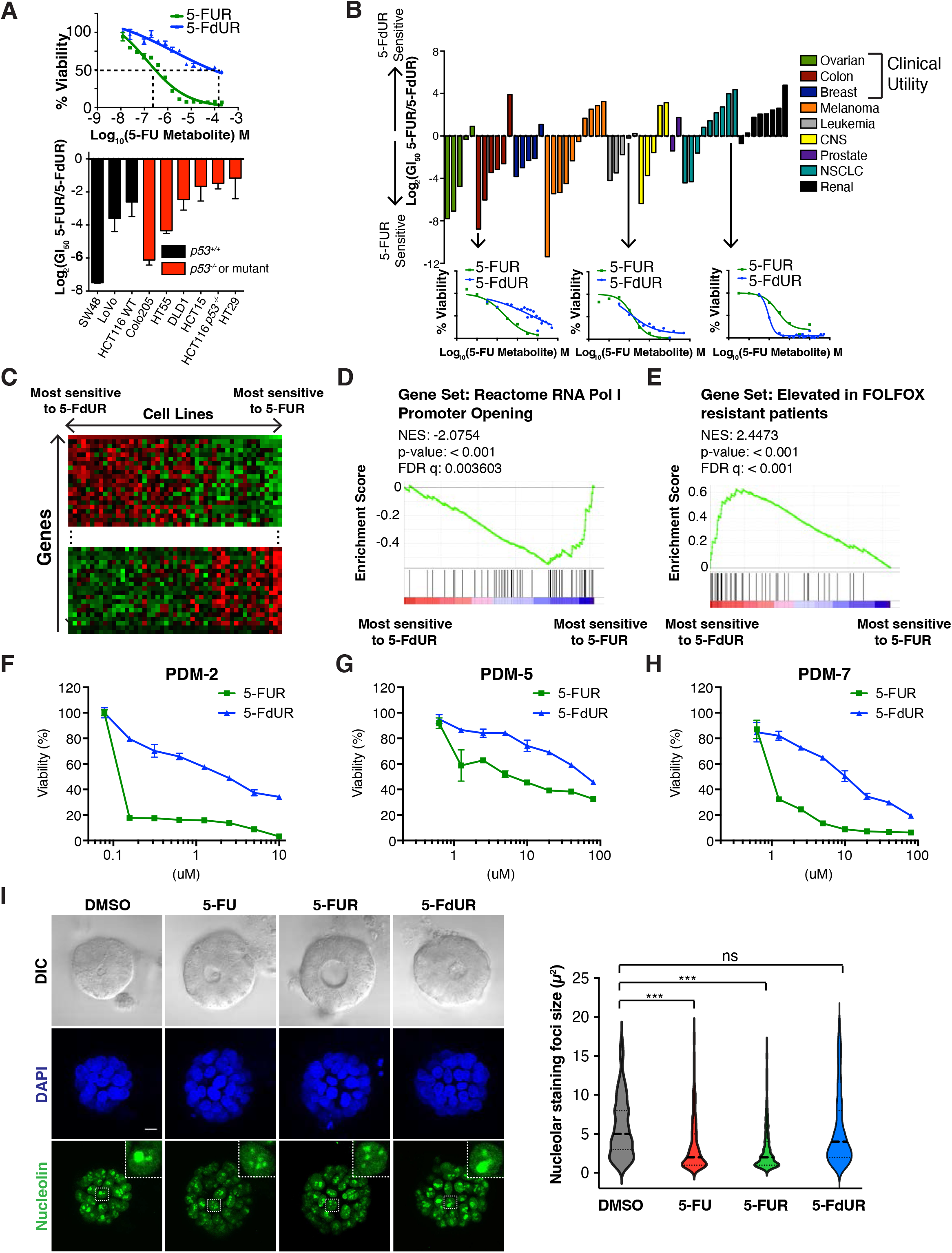
Variance in response to 5-FU metabolites predicts patient response to 5- FUbased therapy. A) A CRC cell line panel was treated with 5-FUR and 5-FdUR dose titrations, and viability was determined after 72 hours by CellTiter-Glo. Shown is Log_2_(GI_50_ 5-FUR/GI_50_ 5-FdUR) to assess the relative sensitivity to 5-FU’s RNA and DNA damage effects. The top panel represented 5-FUR and 5-FdUR dose titrations in SW48 cell line. B) Dose-response curves for 5-FUR and 5-FdUR were calculated for cell lines in the NCI60 from publically available data, and the Log2(GI50 5-FUR/GI50 5-FdUR) was calculated. Each bar represents a single cell line, and bars are colored based on the tissue of origin for the cell line. C) The Z-score of Log_2_(GI_50_ 5-FUR/GI_50_ 5-FdUR) calculated in panel B was correlated with the Z-score of gene expression for the NCI-60 cell line panel to identify genes that were highly correlated with sensitivity to one metabolite or the other. D) The sensitivity-gene expression correlation from panel C was used to identify gene sets that were highly expressed in 5-FUR sensitive and resistant cell lines. Gene sets involved in RNA polymerase I transcription were highly enriched (ES: −0.551, NES: − 2.0754, p value: < 0.001, FDR q value: 0.003603). E) A gene signature that predicts patient sensitivity to FOLFOX suggests that non-responders are resistant to 5-FU mediated RNA damage (ES: 0.6301, NES: 2.4473, p value: < 0.001, FDR q value: < 0.001). F-I) organoid were treated with or without various concentration of 5-FU, 5-FUR and 5- FdUR. Viability was determined using Resazurin cell viability assay after 5 days of drug treatment J) PDM2 organoid were treated with DMSO, 40 μM 5-FU, 1 μM 5-FUR, or 20 μM 5-FUR for 6 hours and subjected to immunofluorescence against nucleolin. The scale bar is 10 μm. The size of nucleolin staining were quantified and plotted as violin plot from more than 100 cells \*\*\**P < 0.001*.

To determine how cells respond to these compounds across different tumor types and tissues of origin, we leveraged publicly available dose-response data from the NCI- 60 cancer cell line panel (Reinhold et al., 2012) and calculated the same relative response metric described above for cell lines in which an accurate GI_50_ could be determined for both compounds (see methods for details). There was substantial variability in the relative sensitivity of cell lines from different tumor types to the RNA and DNA damage effects of 5-FU metabolites across the NCI60 panel, and, surprisingly, it occurred in a tissue-specific manner (Figure 8B). Cell lines from tissues in which 5-FU has shown greatest clinical efficacy—colon, breast, and ovarian (Wilson et al., 2014)—were almost always more sensitive to 5-FUR than 5-FdUR, whereas cells from other tissue types showed greater sensitivity to 5-FdUR or no overall selectivity. These findings indicate that while CRC cells are more sensitive to 5-FUR, this phenomenon is not universally true across cells from other tissues of origin. Further, this suggests that tumor types clinically responsive to 5-FU treatment appear to be particularly sensitive to its RNA damaging effects.

### Gene expression analysis of patient tumor samples suggests that 5-FUR sensitivity is a predictor of clinical response to 5-FU-based therapy

To elucidate patterns of gene expression associated with distinct RNA and DNA damage sensitivity, RNA expression data from the NCI-60 cell line panel was correlated with the metric of relative sensitivity to 5-FUR and 5-FdUR (Figure 8C). Genes were ranked according to their Pearson correlation coefficients and analysed using Gene Set Enrichment Analysis (GSEA) (Subramanian et al., 2005). Importantly, this analysis provided an independent validation of our earlier results by revealing that the expression of genes promoting RNA Polymerase I-dependent transcription across a wide variety of cell lines was strongly enriched amongst genes that correlated with the greatest differential sensitivity to 5-FUR compared to 5-FdUR (Figure 8D and Supplemental Table 1). Furthermore, genes whose expression *decreases* in a 5-FU-*resistant* gastric cancer cell line were found to be highly expressed in our analysis of 5-FUR-sensitive cells (Supplemental Figure 6E), suggesting that resistance to 5-FU is associated with loss of sensitivity to the specific RNA-damaging metabolite 5-FUR.

In contrast, gene sets associated with sensitivity to 5-FdUR were related to cell proliferation and the cell cycle (Supplemental Table 1), consistent with the primary effect of 5-FdUR being mediated through the DDR. Finally, we asked if this data could provide insights into patient responses to 5-FU-based therapies. In an effort to identify gene expression signatures that predict response to FOLFOX, Tsuji et al. took biopsies from patients prior to treatment and determined basal gene expression profiles for each sample (Tsuji et al., 2012). Patients were subsequently classified as responders or non-responders to treatment, allowing those authors to derive a strongly predictive gene signature that was highly expressed in non-responders. We found that their predictive signature for non-responders was highly enriched in genes that anti-correlate with sensitivity to 5-FUR versus 5-FdUR, suggesting that patients whose tumors do not respond to 5-FU-based therapies are likely resistant to 5-FU’s RNA damage effects (Figure 8E).

### 5-FU-induced RNA damage is a more potent inducer of cell death in patient-derived CRC organoids than 5-FU-induced DNA damage

To further explore whether RNA damage is primarily responsible for the dominant cytotoxic effects of 5-FU treatment in colorectal cancer, we directly compared the relative responses of three patient-derived CRC primary tumor organoids to 5-FUR and 5-FdUR. Consistent with our finding in CRC cell lines, 5-FUR was found to be a much more potent inducer of tumor cell death in all three CRC tumor organoids (Figure 8F-H). Examination of the nucleolin, a marker of nucleolar morphology, showed that treatment 5-FU and 5- FUR, but not with 5-FdUR, caused nucleolar restructuring, resulting in the appearance of smaller necklace-like nucleolin foci within 6 hours of treatment (Figure 8I), consistent with impairment of ribosome biogenesis.

## Discussion

Motivated by the unexpected finding of a lack of synergistic, or even additive anti-tumor effects in a systematic study of both human clinical trial data and a panel of colorectal cancer cell lines in response to clinically used 5-FU based combination chemotherapy, we have shown here using systems pharmacology, phospho- and ubiquitin proteomics, cell biology, and biochemical validation that apoptotic cell death induced by 5-FU in colorectal cancer cells is primarily mediated by its effects on ribosomal RNA leading to impaired ribosome biosynthesis, rather than through its effects on DNA replication or direct DNA damage. Importantly, by directly analyzing the differential sensitivity to the RNA and DNA damaging metabolites of 5-FU in the NCI-60 dataset, we observed that this RNA damage-induced death response is particularly over-represented in human cell lines from tumor types where 5-FU has particular clinical utility. In addition, by correlating human gene expression data from patients who responded or failed to respond to 5-FU-based therapies, with the gene expression patterns that dictate the relative sensitivity to 5-FU’s DNA- and RNA-dependent toxicities, we demonstrated that this RNA damage phenomenon appears to control the observed clinical response in colorectal cancer patients treated with 5-FU containing regimens. In agreement with this, we found that the RNA-damaging metabolite of 5-FU was a more potent inducer of cell death in CRC patient-derived organoids than the DNA damaging metabolite, and recapitulated the nucleolar restructuring effects seen when these organoids were treated with the parent compound, 5-FU, consistent with impairment of ribosome biogenesis.

It is important to stress that experiments implicating the RNA incorporation of 5-FU as a significant contributor to cytotoxicity were first performed over 40 years ago (Glazer and Lloyd, 1982; Kufe and Major, 1981). Mechanistic studies in flies and fission yeast have shown that 5-FU can affect RNA processing and the RNA exosome (Mojardin et al., 2013; Silverstein et al., 2011), while large-scale chemogenomic screens and other studies in budding yeast suggested that 5-FU’s mechanism of action could involve an RNA-dependent mechanism (Gustavsson and Ronne, 2008; Lum et al., 2004). Similar RNA processing defects have also been reported in mammalian cells (Burger et al., 2010; Carrico and Glazer, 1979; Doong and Dolnick, 1988; Ghoshal and Jacob, 1994; Greenhalgh and Parish, 1990; Kanamaru et al., 1986; Liang et al., 2021; Samuelsson, 1991; Sun et al., 2007). However, the relative importance of these RNA effects on the drug’s clinical efficacy remains an active area of debate and ongoing investigation, given the ability of 5-FU to activate the DDR, and its well accepted mechanism of TYMS inhibition (Longley et al., 2003; Ludikhuize et al., 2022). The proposed importance of its DNA damage effects in human cells have been highlighted by many studies that have shown that 5-FU’s cytotoxicity can be enhanced by targeting its DNA damage effects (e.g. via TYMS knockdown or Chk1 inhibition or Rad51 inhibition or overexpression of N-methylpurine-DNA glycosylase) (Leguisamo et al., 2017; Muhale et al., 2011; Srinivas et al., 2015; Xiao et al., 2013). Our work, using a broad panel of CRC cell lines, CRC organoids, the NCI-60 collection, RNA and DNA metabolites of 5-FU, and gene expression data from human colorectal cancer patients, indicates that these DNA damage perturbations are likely enhancing the secondary effect of the drug in clinically relevant tumor tissue types, and that 5-FU’s primary efficacy in clinically relevant tumor types is dictated by ribosomal RNA damage and defective ribosome biogenesis, which can be specifically enhanced by modulating rRNA transcription to enhance tumor cell death.

The finding that the RNA Pol I inhibitor CX-5461 markedly suppressed the apoptotic response to 5-FUR, as did the RNA Pol III inhibitor ML-60218, albeit to a somewhat lesser extend (Fig. 3E), indicates that the integrity of structured RNAs (i.e. rRNAs and tRNAs, respectively) is likely what is being monitored by some type of quality control apparatus. It is also important to note that while CX-5461 has been recently described to activate the DDR (Bruno et al., 2020), potentially as a topoisomerase II poison, the doses used in our experiments did not result in activation of the DDR (Figure 2F). Sun et al (Sun et al., 2007) showed that a component of 5-FU-induced apoptosis resulted from stabilization of p53 as a consequence of disrupted rRNA processing, leading to an enhanced interaction of MDM2 with ribosomal proteins. However, given that CX-5461 treatment, which inhibits 5- FU-induced apoptosis, also leads to ribosomal protein-MDM2 interaction, our results indicate that the accumulation of 5-FU-dependent rRNA damage is a more severe form of cellular stress than loss of new rRNA transcription.

The preferential sensitivity to the RNA-damaging effects of 5-FU, rather than the effects of the drug on DNA, is due, in part, to the inherent preference of 5-FU to accumulate in RNA (Figure 2 and (Pettersen et al., 2011), rather than a change in how cells use uracil metabolites during drug treatment. This preferential accumulation is consequential, as we found that RNA damage is a particularly cytotoxic event in CRC cells. Clinical oncology textbooks and recent papers, however, continue to highlight the inhibition of thymidylate synthesis as the major mechanism for the clinical efficacy of 5- FU (Holland-Feei, 2010; Ludikhuize et al., 2022), and direct hepatic artery infusion of the DNA-damaging metabolite of 5-FU, 5-FdUR (Floxuridine™), rather than its RNA-damaging metabolite remains a clinical standard of care for treatment of isolated hepatic metastases from CRC (Doussot et al., 2015). Our findings implicating disruption of ribosome biogenesis accompanied by lysosomal rRNA destruction and ubiquitin-mediated proteolysis of ribosomal proteins as the primary mechanism for 5-FU-induced cell death are strongly supported by a recent study from Nordlund and colleagues (Liang et al., 2021) who reported that RNA modification pathways were strong contributors to the cytotoxicity of 5-FU, but differ from those of Therizols et al., who claimed that 5-FU-treatment resulted instead in the generation of functional 5-FU-containing ribosomes which preferentially translate a set of survival genes (Therizols et al., 2022). The reason for the discrepancy between our data and those of Therizols et al is unclear, although, like us, those authors also observed a marked decrease in the levels of 18S and 28S rRNA after 5-FU treatment, which we showed is accompanied by the disruption of nucleolar morphology and a decrease in total ribosome numbers using cell fractionation and density gradient centrifugation (Figs. 5A and 6A). Furthermore, we showed that treatments that reduced ribosome biosynthesis made cells resistant to the drug, while treatments that enhanced ribosome biosynthesis preferentially sensitized cells to 5-FU. In particular, we showed that activation of rRNA transcription by inhibiting KDM2A significantly enhanced the sensitivity of CRC to the RNA damage effects of 5-FU. Intriguingly, KDM2A is a multi-functional lysine demethylase that is overexpressed in a variety of cancer types, including gastric, colon, breast and non-small cell lung cancer, and thought to contribute to tumor metastasis, in part through Erk1/2 activation (Cao et al., 2018; Chen et al., 2016; Huang et al., 2015; Wagner et al., 2013). Whether its role as a suppressor of rRNA transcription contributes to its pro-metastatic behavior is unclear. Nonetheless, our data clearly implicate inhibition of KDM2A as a potential mechanism to enhance the toxicity of 5-FU in CRC cells.

In the unperturbed cell state ribosome abundance is highly regulated at the level of biogenesis and turnover to meet cellular needs under various physiologically conditions (An and Harper, 2018). Ordinarily, ribosome turnover is regulated entirely through the lysosomal/autophagy pathway (ribophagy), which is enhanced under conditions of nutrient starvation or mTOR inhibition (An and Harper, 2018; Kraft et al., 2008). In contrast, when ribosomal proteins accumulate in stoichiometric excess over the rRNA, (Lam et al., 2007) showed that they were degraded by ubiquitin-dependent proteolysis. Here, we found that 5-FU RNA damage induced a hybrid response in which the fluorinated RNAs were targeted to the lysosome for degradation, while the excess ribosomal proteins were then ubiquitinated and targeted to the proteosome. The molecular mechanism through which this hybrid process activates the apoptotic response remains to be determined.

Importantly, our findings should not be interpreted, in any way, as invalidating the clear clinical utility of FOLFOX and FOLFIRI chemotherapy regimens, both of which have shown marked improvement in outcomes for a selected subset of CRC patients (de Gramont et al., 2000; Saltz et al., 2000), likely as a consequence of patient-to-patient variability (Palmer and Sorger, 2017). Instead, our results reveal the importance of the RNA damage mechanism for 5-FU induced cell death in colon cancer cells, that could be further leveraged by additional drug combinations that specifically target ribosome biogenesis pathways. (Figure 9).

**Figure 9:**
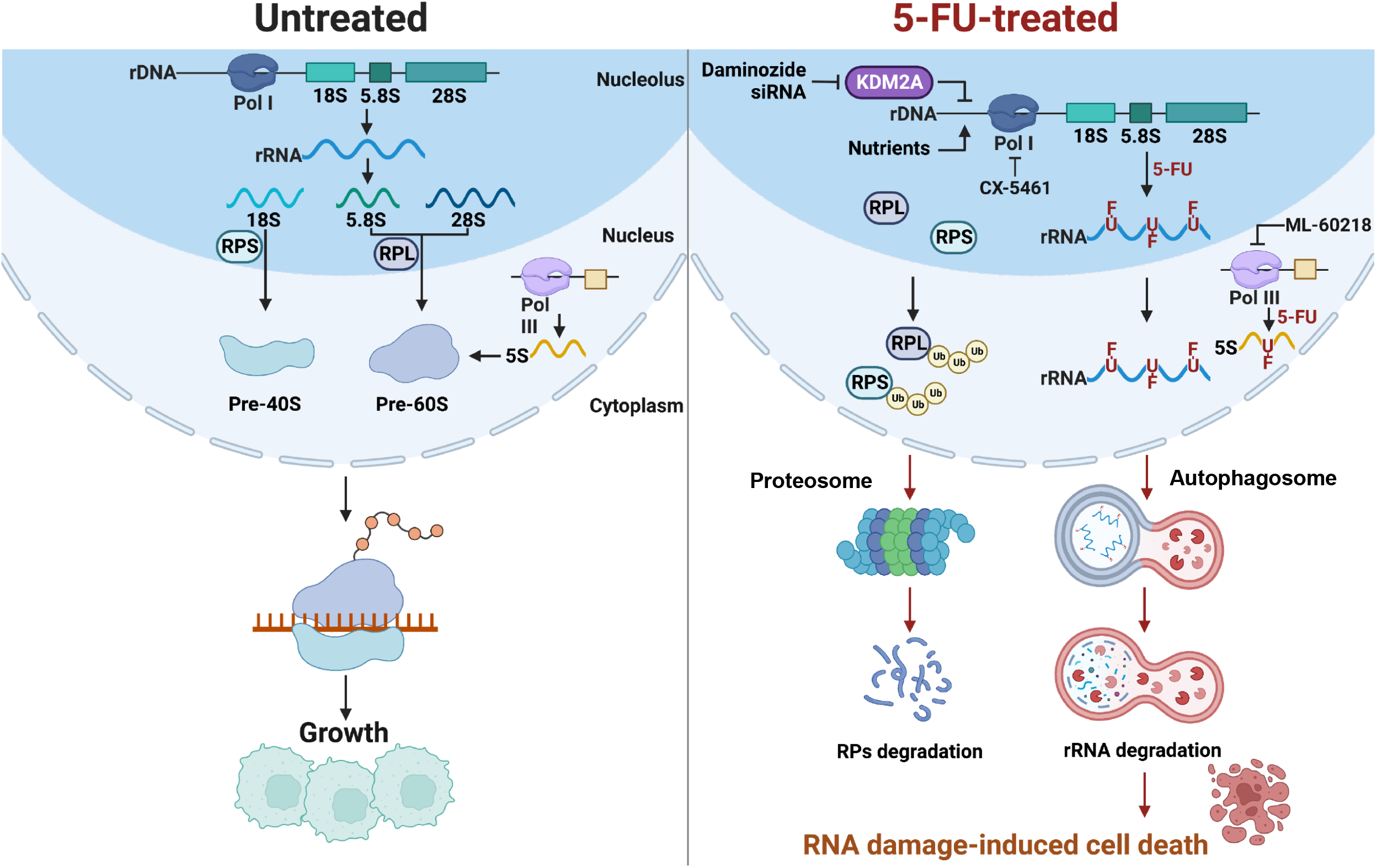
Proposed model for 5-FU-induced RNA damage. 5-FU incorporation into rRNA is proposed to cause rRNA degradation through a autophagosome-mediated pathway. The ubiquitinated ribosome proteins are degraded by proteasome pathway. The 5-FU toxicity can be modulated by targeting the ribosome biogenesis pathway.

## Author Contributions

Conceptualization, J.-K.C., K.A.M., I.G.C., M.B.Y.; Methodology, J.-K.C., K.A.M., J.C.P., Y.W.K., I.G.C., A.M.H., A.I.T., D.S.K., K.Y., C.M.R., G.E., H.H., O.H.Y., A.C.P., M.G.V.H., M.B.Y.; Formal Analysis, J.-K.C., K.A.M., J.C.P., Y.W.K., A.I.T., D.S.K., K.Y., C.M.R., G.E., J.M.H., H.H., M.B.Y.; Investigation, J.-K.C., K.A.M., E.D.H., Y.W.K., L.S.-L., and A.D.; Writing – Original Draft, J.-K.C., K.A.M., and M.B.Y.; Writing – Review & Editing, J.K.C., K.A.M., and M.B.Y.; Funding Acquisition, J.-K.C., K.A.M., M.G.V.H., and M.B.Y.; Supervision, A.C.P., O.H.Y., M.G.V.H., M.B.Y.

## Acknowledgements

We wish to thank Drs. Pau Creixell, Peter Bruno, Michael Hemann, and members of the Yaffe lab for helpful discussions during the course of this project; the Swanson Biotechnology Center at the Koch Institute/MIT for technical support, especially the Flow Cytometry core and High-Throughput Screening core (Jaime Cheah and Christian Soule); Robert Fisher for sharing results prior to publication. Microscopy core (Jeff Wyckoff). Christopher Burge & Gene-Wei Li (James Taggart) lab at MIT for assistance of ribosome profiling. PTMScan is a trademark of Cell Signaling Technology. PTMScan studies are performed under a limited use license from Cell Signaling Technology. This work was supported by the Damon Runyon Cancer Research Fund (DRG 2127-12 to KAM), Ludwig Center at MIT Fellowship (to JKC), NIH grants ES-015339, GM-104047, CA112967 (to MBY), and NIH Cancer Center Support Grant CA014051, the Ovarian Cancer Research Fund, the Charles and Marjorie Holloway Foundation, the STARR Cancer Consortium, and the MIT Center for Precision Cancer Medicine.

## STAR Methods

### Cell culture

HT55, SW48, GP5d, Colo678, Colo320DM, HCT116, and HT-29 cells were grown in DMEM supplemented with 10% FBS and penicillin/streptomycin. HCT15, DLD1, and Colo205 cells were grown in RPMI supplemented with 10% FBS and penicillin/streptomycin. LoVo cells were grown in HAMS F12 supplemented with L-glutamine, 10% FBS and penicillin/streptomycin.

### Immunoblotting

After the indicated treatment, cells were collected with their media, pelleted by centrifugation, and washed 2x PBS. Cell pellets were then lysed in RIPA buffer supplemented with phosphatase inhibitors (PhosSTOP, Roche) and protease inhibitors (cOmplete protease inhibitors, Roche). Lysate concentrations were determined by BCA assay, and equal amounts of protein were loaded in each lane of the SDS-PAGE gel. After electrophoresis, proteins were transferred to nitrocellulose membranes, blocked with Odyssey Blocking Buffer (Licor) diluted 1:1 with PBS. Antibodies were diluted in Odyssey Blocking Buffer mixed 1:1 with PBS-T. Primary antibodies were typically incubated with membranes overnight at 4O C, and secondary antibodies were applied for 1-2 hours at room temperature. After antibody incubations, membranes were washed 3x with PBS-T. Nearly all western blots were visualized Licor Odyssey Imager, and analyzed using Image Studio software (Licor). Western blots in supplemental figure 5B were visualized by chemiluminescence.

### Flow cytometry

After the indicated treatment, cells were collected, fixed, and stained as described in (Lee et al., 2012). Briefly, treated cells were collected by trypsinization and mixed with their growth media to ensure that detached or loosely attached cells were also harvested. Cells were then pelleted by centrifugation, washed with PBS, and fixed with 4% formaldehyde for 15 minutes at room temperature. Cells were washed twice with PBS + 1% BSA, resuspended in ice-cold methanol, and stored at −20° C until further processing. For antibody staining, cells were pelleted by centrifugation, washed twice with PBS-T + 1% BSA, and, if using, incubated with the unconjugated primary antibody for > 1 hour at room temperature while mixing. Fixed cells were then washed twice with PBS-T + 1% BSA, and stained with appropriate secondary antibody and/or a fluorphore-conjugated primary antibody typically overnight at 4° C while mixing. Cells were then washed twice with PBS-T + 1% BSA, once with PBS + 1% BSA. Induction of apoptosis was determined based on positive staining for cleaved-caspase 3 (secondary antibody conjugated with Alexa488) and/or Alexa647-conjugated cleaved-PARP.

### Detection of ^14^C-5-FU and ^14^C-Uracil in RNA and DNA

For cell labeling experiments, 24 hours after cells were plated in a 6-well dish, 0.6 μCi of [2-^14^C]-Uracil (56 mCi/mmol) or [2-^14^C]-5-FU (48 mCi/mmol) was added to each well containing 1 mL of media. Cells were collected at the indicated times. Media was then removed, cells were washed with PBs, and RNA was isolated by TRIzol extraction (Invitrogen), and DNA was isolated from separate wells using the DNeasy Blood and Tissue genomic DNA isolation kit (Qiagen). Incorporation of the compounds was determined by scintillation counting, and counts in each sample were normalized to the input radioactivity for each well.

### siRNA knockdown

For TYMS knockdown, 2.25E6 HCT116 cells were plated in a 10 cm dish, and 24 hours later cells were transfected with 5 nM siControl or siTYMS using Lipofectamine RNAiMAX. 24 hours after transfection, cells were split and 3E5 cells were plated per well in a 6 well plate. Cells were treated as indicated approximately 24 hours later. For ATG5 knockdown, 1E6 HCT116 cells were plated in a 10cm dish, and 24hours later cells were transfected with 10 nM siRNA. 2E6 siRNA-treated cells were plated into 10 cm dish after 24 hours. 24 hours later, cells were treated as indicated. For KDM2A knockdown, 1E6 HCT116 cells were plated in a 10cm dish, and 24hours later cells were transfected with 20 nM siRNA. siRNA-treated cells were plated after 24 hours and treated as indicated after another 24 hours. The ATG5 and KDM2A siRNA sequences were found in GenomeRNAi website and synthesized by Sigma-Aldrich with 3’-UU overhangs.

### Human clinical trials analysis

Kaplan-Meier plots of Progression-Free Survival (PFS) or Time To Progression (TTP) for previously untreated metastatic colorectal cancer treated with 5-FU/LV, oxaliplatin, irinotecan, FOLFOX, and FOLFIRI were extracted and digitized from published clinical trials (Becouarn et al., 1998; Douillard et al., 2000; Giacchetti et al., 2000; Rougier et al., 1997) (Supplemental table 2). The trial of oxaliplatin monotherapy reported median PFS (4.1 months) and median duration of response (7.2 months) among 24% of patients with response (Becouran et al., 1998), which was fitted by a 2-parameter Weibull survival function. Clinical drug additivity was defined as the addition of progression-free survival times (PFS) or times to disease progression (TTP), while correcting for schedules of CT scanning used to measure progression. We used a two-step procedure to simulate the PFS distribution under drug additivity. First, a virtual patient cohort was generated by sampling from a joint distribution of monotherapy responses with a correlation supported by experimental data. Second, each virtual patient’s response to two drugs (A, B) was calculated as the sum of PFS or TTP times for the individual drugs (PFS_AB_ = PFS_A_ + PFS_B_ – first scan time). For example, if a hypothetical patient X had 4 months PFS in response to drug A, and 7 months PFS in response to drug B, and the first CT scan for assessing disease progression was obtained at 1 month, then under drug additivity, the expected PFS time for patient X treated with A + B would be 4 + 7 – 1 = 10 months. The Cox proportional hazard method was used to calculate hazard ratios between the additivity predictions and the observed PFS/TTP for patients treated with the drug combination. Additional details of the methods are described in (Hwangbo et al., 2022).

### Proteome and PTM Quantification Sample Preparation

Lysis buffer containing 9 M urea, 150 mM NaCl, 50 mM HEPES (pH 8.0), one complete-mini (EDTA free) protease inhibitor (Roche), and one phosphatase inhibitor (PHOSstop) tablet (Roche) were added to cell culture plates in order to lyse the cells. Protein concentrations were then determined by Bradford assay and then disulfide bonds were reduced by incubation with 5 mM DTT (45 min, 37°C). This was followed by alkylation of cysteine residues by 15 mM IAA (30 min, RT Dark) which was quenched by the addition of 5 mM DTT (15 min, RT Dark). Initial protein digestion was performed by the addition of LysC (1:50 enzyme:substrate ratio) followed by incubation at 37°C for 3 hours. Samples were then diluted to 1.5 M urea with 50 mM HEPES (pH 8.0) before the addition of Trypsin (1:25 enzyme to substrate ratio) and incubation overnight at 37°C. The next day, the resulting peptide mixtures were acidified and desalted via solid phase extraction (SPE; SepPak, Waters). Following desalting samples were separately enriched for phosphorylated peptides using TiO2 enrichment. Flow through of this enrichment was saved for ubiquitylome and global proteome analysis. Enriched phosphorylated peptides were then desalted and resuspended in 200 mM HEPES (pH 8.5) and labeled with tandem mass tags (TMT, Thermo Fisher Pierce) according to the manufacture instructions. After 1 hour of labeling the reaction was quenched by the addition of 5% hydroxylamine and incubated at room temperature for 15 min. Labeled peptides were then mixed, acidified, and purified by SPE. Labeled and combined phosphopeptides were then enriched for peptides bearing the ATM/ATR motif utilizing a PTMscan (Cell Signaling Technologies) kit. Enriched ATM/ATR peptide were desalted before nLC-MS/MS analysis. Flow through from the ATM/ATR enrichment was desalted before separation into 96 fractions by offline basic-reversed phase chromatography. Fractions were then combined into 12 fraction pools which were analyzed by nLC-MS/MS. For ubiquitylome and global proteome analysis flow through from the TiO2 enrichment was desalted before enrichment with the PTMscan Kgg remnant motif antibody (Cell Signaling Technology) as previously described (Rose et al., 2016). This resulted in 6 Kgg fractions which were desalted before nLC-MS/MS analysis. Flow through from the Kgg enrichment was was desalted before separation into 96 fractions by offline basic-reversed phase chromatography. Fractions were then combined into 24 fraction pools of which 12 were analyzed by nLC-MS/MS.

### Proteome and PTM Quantitative nLC-MS/MS Analysis

For nLC-MS/MS analysis, peptides were separated using a Dionex UltiMate 3000 RSLCnano Proflow system (Thermo Fisher Scientific) for ATM/ATR and Kgg experiments or a NanoAcquity UPLC (Waters) for global proteome and phosphorylation experiments. A gradient of 2% buffer A (98% H20, 2% ACN with 0.1% formic acid) to 30% or 35% buffer B (98% ACN, 2% H20, 0.1% formic acid) with a flow rate of 450 or 500 nL/min was used to separate peptides over a 25 cm capillary column (100 µm I.D.) packed with Waters nanoAcquity M-Class BEH (1.7 µm) material (New Objective, Woburn, MA). Samples were analyzed using an Orbitrap Fusion or Orbitrap Fusion Lumos mass spectrometer, (Thermo Fisher Scientific, San Jose, CA). For all analyses the SPS-MS3 method was implemented for improved quantitative accuracy (McAlister et al., 2014; Ting et al., 2011). For all experiments, intact peptides were surveyed in the Orbitrap and the top 10 peptides were selected for fragmentation. For Kgg experiments +2 precursor ions were not selected for fragmentation as previously described.^0^ For global proteome, global phosphoproteome, and ATM/ATR experiments fragments were analyzed in the ion trap while Kgg experiments utilized the Orbitrap for MS^2^ analysis. Quantitative MS^3^ scans selected the 8 most abundant fragment ions from the MS^2^ spectrum and fragmented them at high energy (HCD, 55 NCE) to produce reporter mass ions. All raw data can be found within the MassIVE repository with the identifier: MSV000090256 (reviewer login = MSV000090256_reviewer & password = multiome).

### Proteomic and PTM Data Analysis

Assignment of MS/MS spectra was performed using the MASCOT search algorithm to search against all entries for *Homo sapiens* (human) in UniProt (downloaded June 2016). A search of all tryptic peptides (2 missed cleavages) was performed and a precursor tolerance of 50 ppm was used to limit the number of candidate peptides, while a 0.8 Da tolerance was used to match MS/MS data collected in the ion trap. Static modifications included TMT on the N-terminus of peptides and lysine residues (+229.16293 for TMT) and cysteine alkylation (+57.0215), while variable modifications included methionine oxidation (+15.9949) for all experiments, phosphorylation of S/T/Y (+79.9663) for global phosphorylation and ATM/ATR experiments, and gg remnant (+114.0429) for ubiquitylome experiments. Peptide spectral matches were filtered to a 2% false discovery rate using a target decoy approach scored with a linear discrimination analysis algorithm before filtering to a 2% false discovery rate at the protein level as previously described (Kirkpatrick et al., 2013). Quantitative values were extracted and corrected for isotopic impurities using Mojave (Zhuang et al., 2013). Additionally, quantitative events with a precursor purity < 0.5 (± 0.25 Da) or sum intensity < 30,000 were discarded before quantitative values were normalized and converted to “relative abundance” values using custom scripts coded in R. For quantitative analysis, peptide spectral match level data was summed to the peptide level and peptide level data was summed to the protein level. For each comparison a Student’s t-test (two tailed, unequal variance) was used to calculate a nominal p-value. Data were further visualized in Spotfire (Perkin Elmer, Waltham, MA).

### Computational Analysis of NCI-60 Cell Line Drug Response

Dose-response data for 5-FUR and 5-FdUR treatment in the NCI-60 cell line panel was downloaded from the NCI’s Developmental Therapeutics Program website. Percent growth inhibition values were averaged when multiple measurements were reported for a given compound concentration in the cell line being examined. Dose response curves were fit using Prism, and absolute GI_50_ measurements were extracted for each cell line treatment. Quality of the curve fit was determined R^2^ > 0.70, and cell lines in which either compound did not fit this criteria were excluded from further analysis. Further, if a GI_50_ could not be accurately determined for a compound due to the GI_50_ being outside of the reported dose range (i.e. the lowest concentration of a compound gave greater than 50% inhibition or the highest concentration gave less than 50% inhibition), then this cell line was also excluded from further analysis. To assess relative sensitivity of the remaining cell lines to 5-FUR and 5-FdUR, we calculated the log2(5-FUR GI_50_/5-FdUR GI_50_).

For the cell lines that passed our data quality filters above, we downloaded gene expression data collected by microarray from the NCI CellMiner website. The Z score of both gene expression data and log_2_(5-FUR GI_50_/5-FdUR GI_50_) were calculated, and the Pearson correlation coefficient between the expression of each gene and the relative sensitivity metric was calculated using MeV. These correlation coefficients were then used as a pre-ranked list for Gene Set Enrichment Analysis (GSEA). Reported statistics are based on gene list permutations.

### Cell survival assays

1000 colorectal cancer cells were plated per well in 96 or 384-well plates. 24 hours after plating drugs were added to the media at the indicated concentrations, and viability was measured 48 or 72 hours after drug addition by either CellTiter-Glo, Resazurin (Figure 1A, B, and F and Supplemental Figure 1 only as indicated. Percent viability is normalized to vehicle treated control wells. Dose-response curves were fit using Prism, and the area under the curve (AUC) and GI_50_ values for response to the compounds were determined when indicated. For dose response matrices, the expected viability was determined based on a Bliss Independence model of drug interaction (Greco et al., 1995), and relative synergy was determined based on comparing the expected viability to that which was observed. For Trypan Blue exclusion viability test, cell sample was diluted 1:1 with 0.4% Trypan Blue solution and incubated for 1-2 minutes at room temperature. The unstained cells were then counted under light microscope.

### Colony formation assay

HCT116 cells treated with different siRNA were counted and plated in a six-well plate. Cells were exposed to different dosages of drugs and then incubated at 37 °C for 10 to 14 days until colonies could be visualized under a light microscope. The colonies were stained with 20% ethanol solution containing 0.5% crystal violet. After gentle washing, the colonies were counted and normalized to control siRNA-treated cells.

### Immunofluorescence

Cells seeded on poly-L-lysine coverslips (BD Biosciences) were fixed with 10% neutral buffered formalin solution (Sigma) for 15 min and extracted with PBS containing 0.5% Triton X-100 (Sigma) for 10 min at room temperature. After blocking with 5% BSA (Sigma), samples were incubated with indicated primary antibodies overnight at 4 °C. Samples were then washed and incubated with secondary antibodies plus DAPI (Thermo Scientific) for 1 h at room temperature. Samples were mounted onto glass slides with Fluoromount mounting medium (SouthernBiotech) and visualized by FV1200 (Olympus) confocal microscope. For live cell microscopy, cells seeded on 35 mm glass-bottom dishes (In Vitro Scientific) were stained with SYTO^TM^ RNA select^TM^ green fluorescence (ThermoFisher) for 20 min at 37 °C, washed and followed with 50 μM monodansylcadaverine (Sigma) plus Hoechst33342 (Invitrogen) for 30 min at 37 °C. After wash with PBS, the stained cells in FluoroBrite DMEM (ThermoFisher) were analysed with FV1200 (Olympus) confocal microscope.

### RNA extraction, cDNA synthesis and Quantitative PCR

Total RNA was extracted using RNeasy mini kit (Qiagene). The RNA concentration was determined using a Nanodrop ND1000 (Nanodrop Technologies). RNA (1 μg) was reversely transcribed to cDNA with SuperScript® III Reverse Transcriptase (Invitrogen) using Random hexamer (Invitrogen) as primer. Real-time PCR was performed on a StepOnePlus instrument (Applied Biosystems) using SYBR^®^ green Master Mix (Applied Biosystems). The qPCR conditions were as followed: 95 °C for 10 min, 40 cycles of 95 °C for 15 s, 60 °C for 1 min. This was followed by melting curve analysis to confirm singlely amplified product: 95 °C for 15 s, 60 °C for 1 min, 0.05 °C/s increment from 60 °C to 95 °C, and hold at 95 °C for 15 s. The results were analyzed with StepOne real-time PCR software.

### Ribosome profiling

Sucrose density gradient centrifugation was used to separate ribosomes into polysomal and subpolysomal fractions. Cells were pre-treated with 0.1 mg/ml cycloheximide for 5 min at 37°C. After washed with PBS containing 0.1 mg/ml cycloheximide, Cells were lysed with lysis buffer (15 mM Tris-HCl (pH 7.5), 0.3M NaCl, 15 mM MgCl_2_, 1 mg/ml heparin, 1% Triton X-100, 0.1 mg/ml Cycloheximide and 80U/ml RNase inhibitor) according to cell number (1.5×10^7^/ml). After centrifugation at 13,000 g for 15 min, the supernatants were loaded onto a 10∼60% sucrose gradients in lysis buffer without Triton X-100. The gradients were sedimented at 38,000 rpm for 2 hours using a SW41 Ti rotor at 4°C (221,777 g). Fractions were collected and analysed with a Gradient Master^TM^ fraction collector system (Biocamp).

### 18S/28S rRNA detection

Total RNA was extracted from the same number of cells using RNeasy mini kit (Qiagene). The RNA was separated using standard formaldehyde denaturing gel for northern blot. Gel was stained with GelRed Nucleic Acid Staining dye (Biotium) and visualized under UV light. 18S and 28S rRNA were quantified by Fiji ImageJ software.

### Organoid culture and drug responses assay

PDM2, PDM5 and PDM7 large intestine organoids were purchased from ATCC. Media for the organoids is composed of advanced DMEM:F12, 100 U/mL of penicillin (Invitrogen, USA), 100 µg/mL of streptomycin (Invitrogen, USA), and 0.25 µg/mL amphotericin (Invitrogen, USA). The following compounds were also added to the growth media: B27(Invitrogen, USA), 10 uM nicotinamide (Sigma-Aldrich, USA), 50 ng/mL EGF (Novus Biologicals, USA), 500 nM A83-01 (Cayman Chemical, USA), 10 uM SB202190 (Cayman Chemical, USA), and 500 nM PGE2 (Cayman Chemical, USA). The organoids were propagated once every week, using standard manual dissociation of Matrigel (Corning, USA), centrifugation at 200 x g for 2 min, and brief trypsinization for 5 min, upon which the cells were re-embedded in Matrigel at a 1:4 passaging dilution. For drug dose response assays, organoids were dissociated to singles cells and plated in Matrigel plugs 10 microliters per well, each containing 2000 cells. Cell number was quantified using resazurin reagent (Sigma-Aldrich, USA) (0.05 mg/ml final concentration) applied to the organoid plugs at endpoint. The organoids were incubated with resazurin for 4 hours, and red fluorescence was quantified by a plate reader spectrophotometer 560ex/590em (Tecan, Männedorf, Switzerland).

### Organoid immunofluorescence

Organoids were washed once with PBS and then fixed by 4% paraformaldehyde for 20 min at RT and stored in PBS at 4°C. Organoids were permeabilized with PBS/3% BSA containing 1% Triton X-100 for 4h at 4°C and then stained with primary antibodies (nucleolin 1:200, cell signaling) overnight in PBS/3% BSA solution. After wash with PBS for 10 min x3, organoids were stained with secondary Ab overnight in PBS/3% BSA and then with 5 µg/mL DAPI (Thermo Scientific) in PBS for 1 hour at 4°C. Imaging was performed using an Olympus FV1200 confocal microscope.

## Key Resource Table

| REAGENT or RESOURCE | SOURCE | IDENTIFIER |
| --- | --- | --- |
| **Antibodies** |  |  |
| Chk2 T68-P | Cell Signaling Thechology | 2661 |
| Chk1 S345-P | Cell Signaling Thechology | 2348 |
| TYMS | Cell Signaling Thechology | 9045 |
| PARP | Cell Signaling Thechology | 9542 |
| cleaved-PARP | Cell Signaling Thechology | 5625 |
| S6 S240/244-P | Cell Signaling Thechology | 2215 |
| Actin | Cell Signaling Thechology | 4967 |
| eIF2a S51-P | Cell Signaling Thechology | 3398 |
| ATF4 | Cell Signaling Thechology | 11815 |
| Actin | Sigma-Aldrich | A-5441 |
| Vinculin | Abcam | ab18058 |
| p53 | Santa Cruz Biotechnology | sc-126 |
| eIF2a | Santa Cruz Biotechnology | sc-133132 |
| cleaved-PARP | BD Biosciences | 558710 |
| cleaved-caspase 3 | BD Biosciences | 559565 |
| RPL26 | Bethyl Laboratories | A300-686A-T |
| RPL7 | Bethyl Laboratories | A300-741A-T |
| RPS15A | Bethyl Laboratories | A304-990A-T |
| ATG5 | Cell Signaling | 12994S |
| KDM2A | Novus | NB100-74602 |
| Nucleolin | Cell Signaling | 14574 |
| **Chemicals** |  |  |
| 5-fluorouracil (5-FU) | Sigma-Aldrich | F6627 |
| 5-fluorouridine (5-FUR) | Sigma-Aldrich | F5130 |
| 5-fluoro-2'-deoxyuridine (5-FdUR) | Sigma-Aldrich | F0503 |
| 5-fluoro-5'-deoxyuridine (5-F-5'dUR) | Sigma-Aldrich | F8791 |
| Aphidicolin | Sigma-Aldrich | 89458 |
| Hydroxyurea | Sigma-Aldrich | H8627 |
| ISRIB | Sigma-Aldrich | SML0843 |
| Irinotecan | Sigma-Aldrich | I406 |
| CX-5461 | Selleck Chemicals | S2684 |
| Oxaliplatin | Selleck Chemicals | S1224 |
| PF-477736 | Selleck Chemicals | S2904 |
| ML-60218 | EMD Millipore | 557043 |
| [2-<sup>14</sup>C]-Uracil | Moravek Biochemicals | MC-124 |
| [2-<sup>14</sup>C]-5-FU | Moravek Biochemicals | MC-101 |
| Chloroquine diphosphate | SIGMA | C6628 |
| MG132 | Selleckchem | S2619 |
| Heparin Sodium Salt | SIGMA | H3149-25ku |
| Monodansylcadaverine (MDC) | SIGMA | D4008 |
| Daminozide | Selleckchem | S4800 |
| Critical Commercial Assays |  |  |
| CellTiter-Glo | Promega | G7572 |
| Resazurin | Biotium | 30025 |
| Lipofectamine RNAiMAX | Invitrogen | 13778075 |
| SYTO60 | Invitrogen | S11342 |
| SYTO<sup>TM</sup> RNA Select | ThermoFisher | S32703 |
| Click-iT<sup>TM</sup> RNA Alexa Fluor<sup>TM</sup> 488 | ThermoFisher | C10329 |
| Experimental Models: Cell Lines |  |  |
| Human: HCT116 WT | ATCC | CCL-247 |
| Human: HT29 | ATCC | HTB-28 |
| Human: Colo320DM | ATCC | CCL-220 |
| Human: Colo678 | K. Haigis (Beth Israel Medical Deconness) |  |
| Human: GP5d | K. Haigis (Beth Israel Medical Deconness) |  |
| Human: LoVo | K. Haigis (Beth Israel Medical Deconness) |  |
| Human: Colo205 | K. Haigis (Beth Israel Medical Deconness) |  |
| Human: HT55 | K. Haigis (Beth Israel Medical Deconness) |  |
| Human: DLD1 | K. Haigis (Beth Israel Medical Deconness) |  |
| Human: HCT15 | K. Haigis (Beth Israel Medical Deconness) |  |
| Human: SW48 | K. Haigis (Beth Israel Medical Deconness) |  |
| Human: HCT116 p53-/- | K. Haigis (Beth Israel Medical Deconness) |  |
| Experimental Models: Organoids |  |  |
| Human: PDM2 | ATCC | HCM-CSHL-0057-C18 |
| Human: PDM5 | ATCC | HCM-CSHL-0061-C18 |
| Human: PDM7 | ATCC | HCM-CSHL-0063-C18 |
| Human: PDM9 | ATCC | HCM-CSHL-0065-C20 |
| Oligonucleotides |  |  |
| SilencerSelect Negative Control No. 1 siRNA | Invitrogen | 4390844 |
| TYMS SilencerSelect siRNA | Invitrogen | s14538 |
| On-TARGETplus non-targeting pool | Dharmacon | D-001810-10-05 |
| ATG5 siRNA-1 | GenomeRNAi, synthesized by Sigma-Aldrich | s18158 |
| ATG5 siRNA-2 | GenomeRNAi, synthesized by Sigma-Aldrich | s18159 |
| ATG5 siRNA-3 | GenomeRNAi, synthesized by Sigma-Aldrich | s18160 |
| KDM2A siRNA-1 | GenomeRNAi, synthesized by Sigma-Aldrich | s22790 |
| KDM2A siRNA-2 | GenomeRNAi, synthesized by Sigma-Aldrich | s22792 |
| Software and Algorithms |  |  |
| Multiple Expression Viewer (MeV) |  | http://mev.tm4.org/ |
| Gene Set Enrichment Analysis (GSEA) | Subramanian, et al.<br>PNAS (2005) | http://software.broadinstitute.org/gsea/index.jsp |
| Other |  |  |
| NCI60 Drug Response and<br>Gene Expression Datasets | Reinhold et al, Cancer Research (2012) | https://dtp.cancer.gov/;<br>https://discover.nci.nih.gov/cellminer/home.do |

**Supplemental Figure 1:**
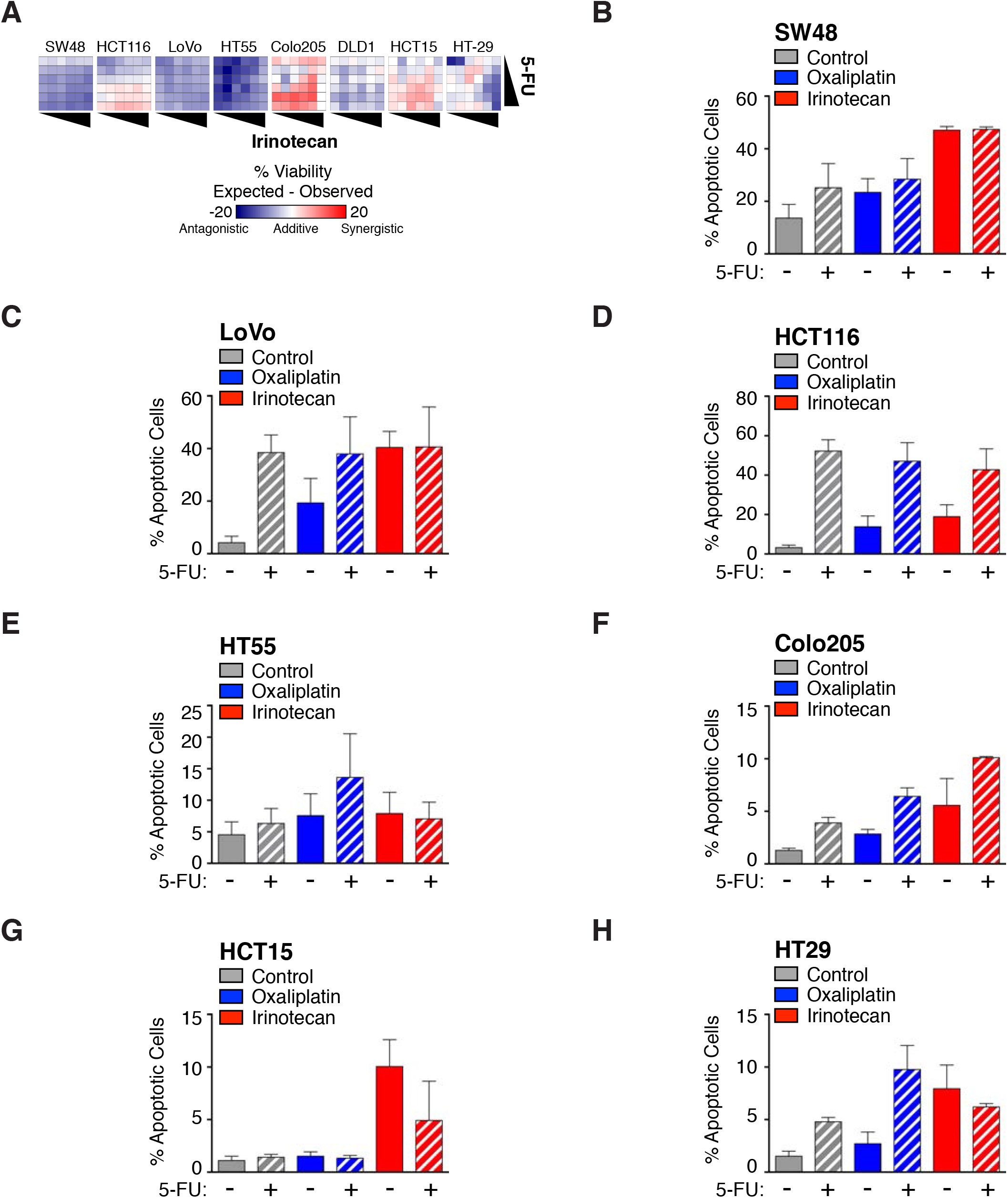
A) Dose-response matrices for the indicated cell lines were generated with 5-FU and irinotecan (top doses of 200 μM and 25 μM, respectively), and viability was measured using Resazurin cell viability assay after 72 h of drug treatment. B) SW48 cells were treated with 5 μM oxaliplatin, 6.25 μM irinotecan +/- 50 μM 5-FU for 48 hours. Cells were collected and fixed for flow cytometry to determine the percentage of apoptotic cells based on c-PARP and c-caspase 3 positivity. C) As in A), except using LoVo cells. D) As in A), except using HCT116 cells. E) As in A), except using HT55 cells. F) As in A), except using Colo205 cells. G) As in A), except using HCT15 cells. H) As in A), except using HT-29 cells.

**Supplemental Figure 2:**
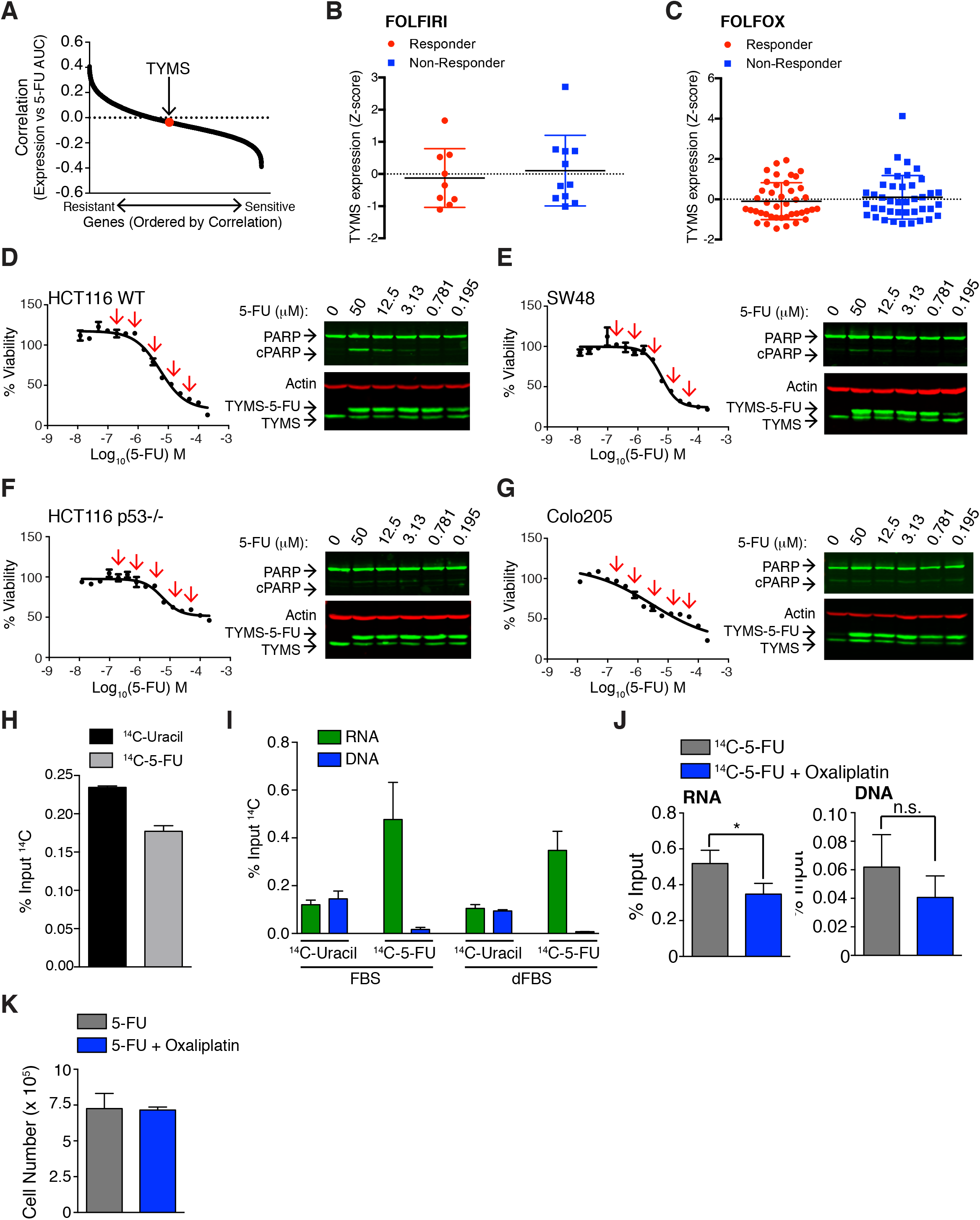
A) mRNA expression was correlated with the area under the curve (AUC) of response for 5-FU for the CCLE cell line panel. In this dataset, TYMS expression poorly correlated with sensitivity to 5-FU. Original drug response data is from Rees MG, et al, 2016. B) TYMS mRNA expression was compared between CRC patients who were responders or non-responders to FOLFIRI chemotherapy. Original drug response data is from Del Rio et al., 2007. C) TYMS mRNA expression was compared between CRC patients who were responders or non-responders to to FOLFOX chemotherapy. Original drug response data is from Tsuji et al., 2012. D) Response to 5-FU in HCT116 WT cells was measured by CellTiter-Glo 72 hours after drug treatment (left). The doses of 5-FU indicated with red arrows were used to treat cells for 24 hours, and acute effects of the drug were measured by PARP cleavage and formation of the inhibited TYMS-5-FU complex. Doses of 5-FU that cause maximal inhibition of the enzyme are not sufficient to cause significant loss of viability in either assay. E) As in D), except in SW48 cells. F) As in D), except in HCT116 *p53*^-/-^ cells. G) As in D), except in Colo205 cells. H) There is a minor difference in the initial uptake of ^14^C-5-FU and ^14^C-Uracil by HCT116 cells. Cells were treated with the radiolabeled compounds for 30 minutes, then collected, washed, and total cellular uptake was measured by scintillation counting. I) The difference in distribution of uracil and 5-FU is maintained when cells are grown in media containing dialyzed serum (dFBS). J) HCT116 cells were treated for 24 hours with ^14^C-5-FU +/- 5 μM oxaliplatin, total RNA and DNA were isolated, and the amount of drug in RNA and DNA was determined by scintillation counting. K) HCT116 cells were treated with 12.5 μM 5-FU (the equivalent dose used in F) and G)) +/- 5 μM oxaliplatin, and cell number was determined.

**Supplemental Figure 3:**
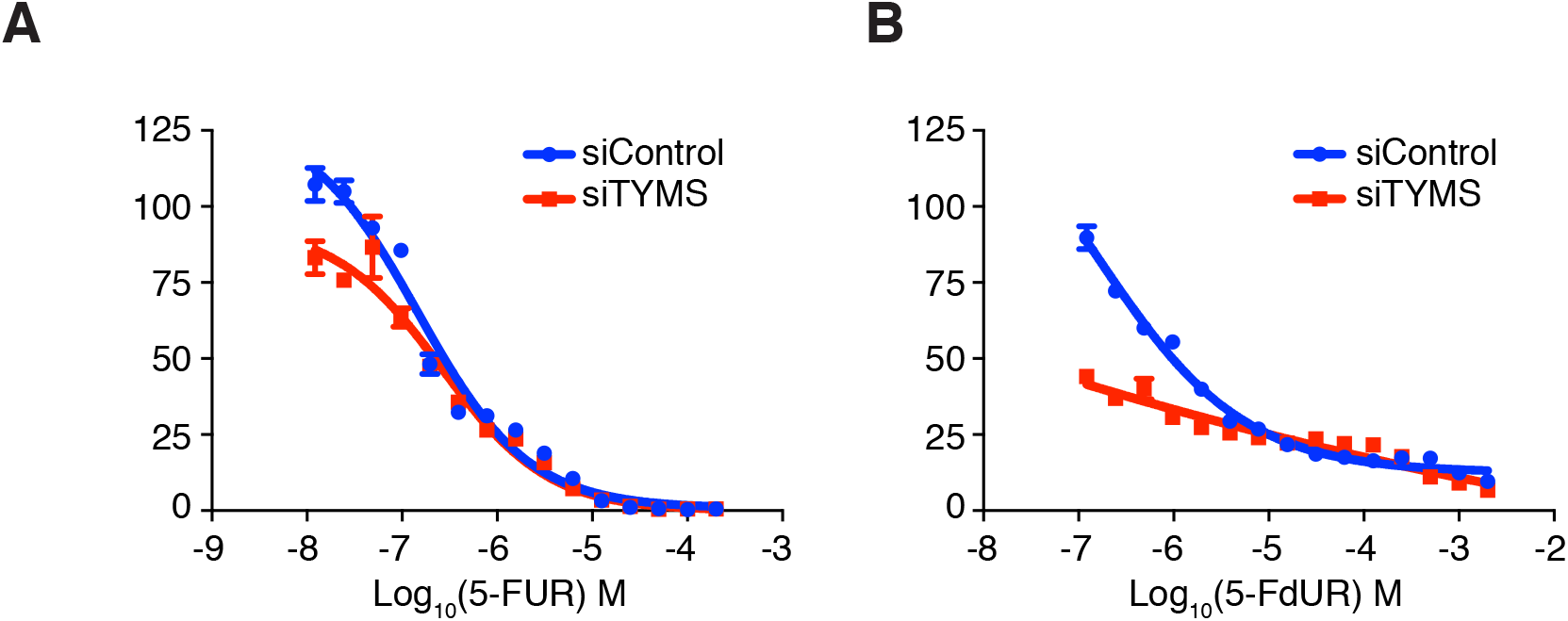
A) HCT116 cells were transfected with siControl or siTYMS for 48 hours, and treated with the indicated concentrations of 5-FUR or 5-FdUR. Viability was determined after 72 hours by CellTiter-Glo.

**Supplemental Figure 4:**
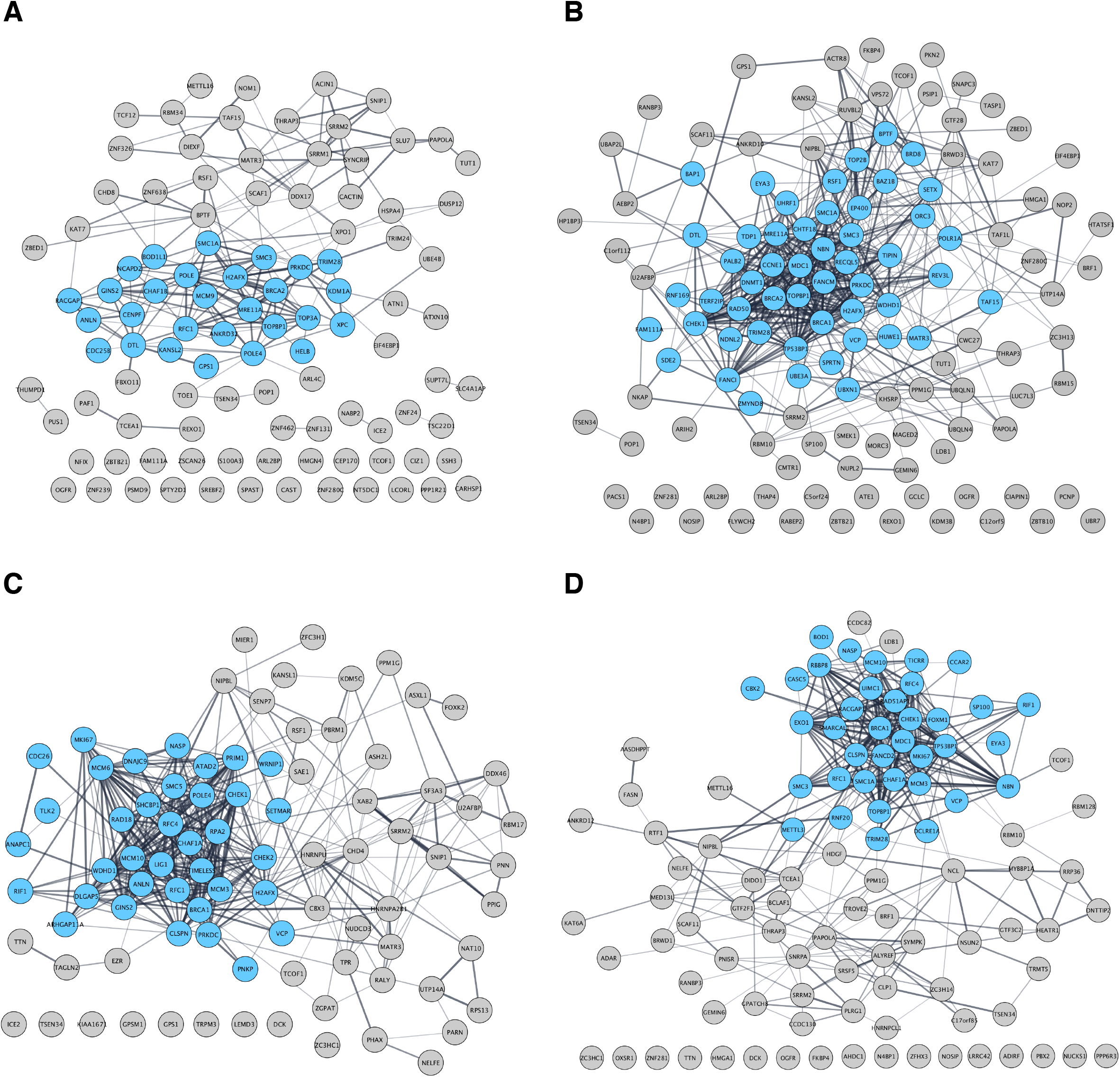
A) STRING functional protein association networks of up-regulated 5-FdUR specific p- SQ/p-TQ proteins. B) STRING functional protein association networks of up-regulated 5-FdUR > 5-FUR p- SQ/p-TQ proteins. C) STRING functional protein association networks of up-regulated 5-FdUR specific total phosphorylated proteins. D) STRING functional protein association networks of up-regulated 5-FdUR > 5-FUR total phosphorylated proteins. The data were generated by Cytoscape software and the proteins involving in DNA repair & DNA replication were labelled in blue.

**Supplemental Figure 5:**
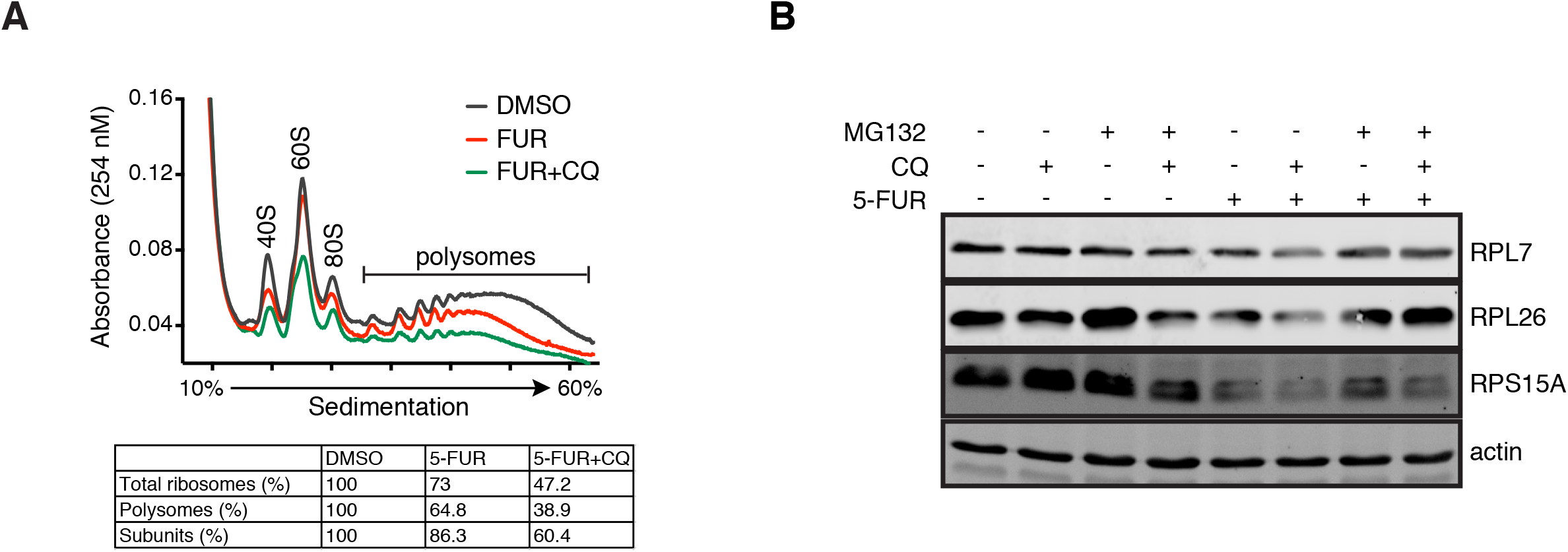
A) HCT116 cells were treated with 2 μM 5-FUR or 2 μM 5-FUR plus 20 μM chloroquine for 18 hours and subjected to polysome gradient to reveal ribosome profile. B) Representative western blot results of Figure 5C.

**Supplemental Figure 6:**
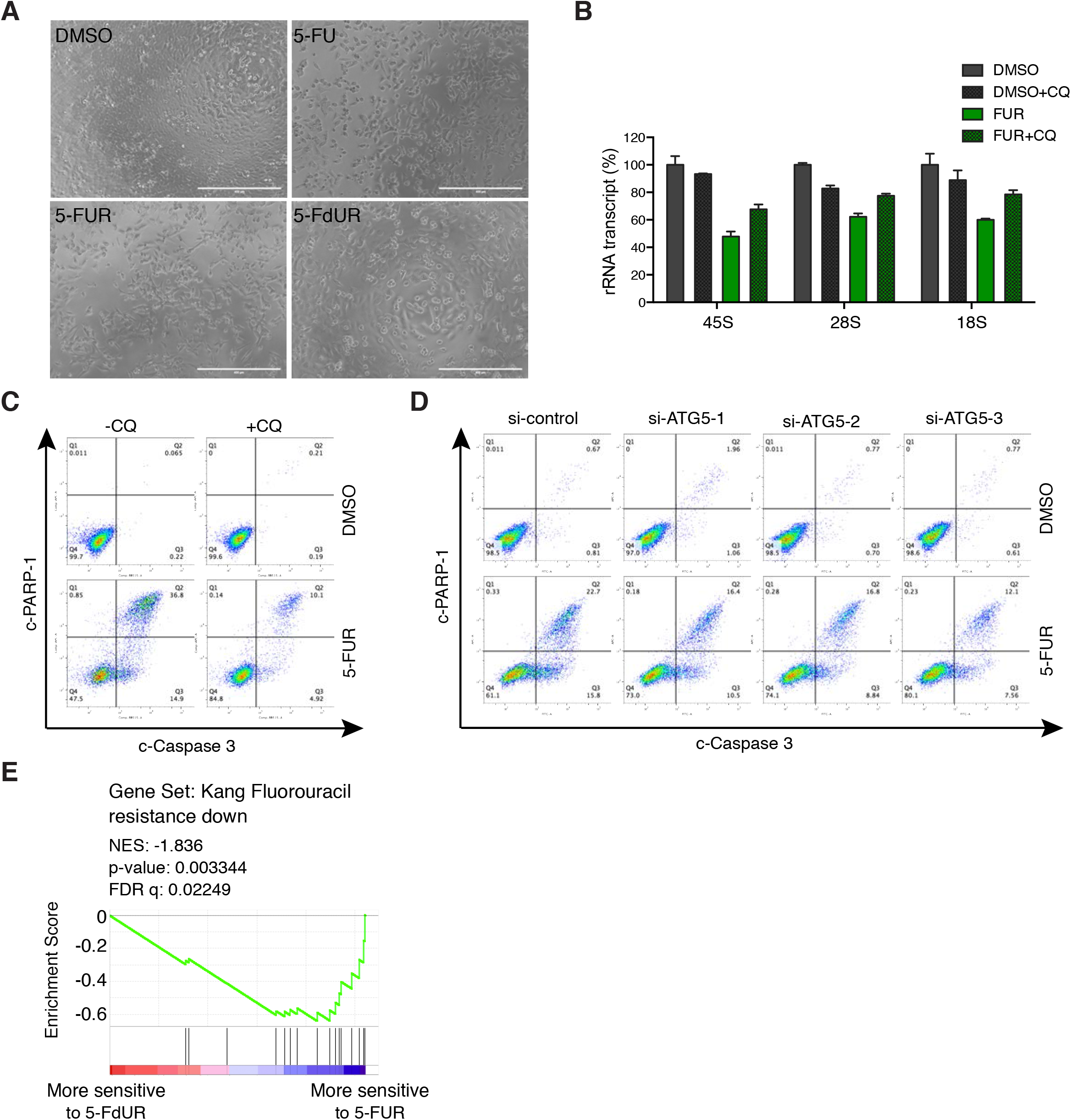
A) Cell morphology of DMSO, 40 μM 5-FU, 2 μM 5-FUR and 2 μM 5-FdUR-treated cells. B) HCT116 cell were treated with or without 20 μM chloroquine and 2 μM chloroquine for 18 hr, ribosomal RNAs were analysed by qPCR (representative of three independent experiments). C) Representative FACS data for Figure 6G. D) Representative FACS data for Figure 6I. E) Cell lines that are more sensitive to 5-FUR highly express genes whose expression decreases in a gastric cancer cell line that acquires resistance to 5-FU (ES: −0.6409, NES: −1.836, p value: 0.003344, FDR q value: 0.02249).

**Supplemental Figure 7:**
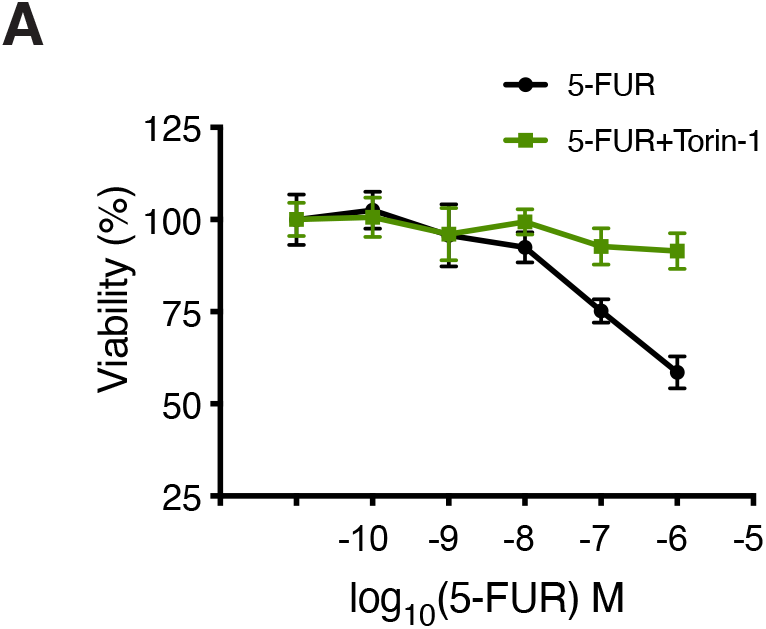
HCT116 cells were treated increasing concentration of 5-FUR +/- 0.3 mM Torin-1 for 24 hours and viability was assessed by MTT assay. (representative of three independent experiments).

**Supplemental Table 1:** Reactome gene sets that are significantly enriched in cell lines with greatest differential sensitivity to 5-FUR and 5-FdUR. To characterize which pathways might affect relative response to 5-FUR and 5-FdUR, we identified significantly enriched Reactome gene sets (FDR q-value < 0.05) for the correlation between gene expression and 5-fluoropyrimidine sensitivity shown in Figure 4C.

| Gene Set | ES | NES | Nom p-value | FDR q-value |
| --- | --- | --- | --- | --- |
| Gene sets enriched in 5-FUR-sensitive cells |  |  |  |  |
| RESPIRATORY ELECTRON TRANSPORT ATP SYNTHESIS BY CHEMIOSMOTIC COUPLING AND HEAT PRODUCTION BY UNCOUPLING PROTEINS | -0.52478445 | -2.1668122 | < 0.001 | 0.0010044 |
| TCA CYCLE AND RESPIRATORY ELECTRON TRANSPORT | -0.48485428 | -2.1210828 | < 0.001 | 0.001962681 |
| RNA POL I PROMOTER OPENING | -0.55097204 | -2.0754452 | < 0.001 | 0.00360292 |
| PACKAGING OF TELOMERE ENDS | -0.5605999 | -2.050865 | < 0.001 | 0.00456106 |
| RESPIRATORY ELECTRON TRANSPORT | -0.514806 | -2.0495539 | < 0.001 | 0.004385635 |
| TELOMERE MAINTENANCE | -0.47566223 | -1.8806362 | < 0.001 | 0.023079738 |
| AMINO ACID SYNTHESIS AND INTERCONVERSION TRANSAMINATION | -0.6399725 | -1.8789145 | < 0.001 | 0.023195667 |
| AMYLOIDS | -0.47619024 | -1.872435 | < 0.001 | 0.023922522 |
| RNA POL I TRANSCRIPTION | -0.46100852 | -1.8541875 | < 0.001 | 0.024723269 |
| BASE EXCISION REPAIR | -0.6184477 | -1.8459262 | 0.001692047 | 0.026708458 |
| RESOLUTION OF AP SITES VIA THE MULTIPLE NUCLEOTIDE PATCH REPLACEMENT PATHWAY | -0.6267394 | -1.8217123 | 0.001845019 | 0.032496147 |
| Gene sets enriched in 5-FdUR-sensitive cells |  |  |  |  |
| THROMBIN SIGNALLING THROUGH PROTEINASE ACTIVATED RECEPTORS PARS | 0.5900197 | 2.1296227 | < 0.001 | 0.002091926 |
| LOSS OF NLP FROM MITOTIC CENTROSOMES | 0.5067976 | 2.1197846 | < 0.001 | 0.002164982 |
| MITOTIC PROMETAPHASE | 0.47416812 | 2.1126013 | < 0.001 | 0.002297129 |
| APC C CDH1 MEDIATED DEGRADATION OF CDC20 AND OTHER APC C CDH1 TARGETED PROTEINS IN LATE MITOSIS EARLY G1 | 0.4841238 | 2.0636468 | < 0.001 | 0.003513513 |
| G ALPHA Z SIGNALLING EVENTS | 0.5301024 | 2.0489228 | < 0.001 | 0.003953945 |
| VIF MEDIATED DEGRADATION OF APOBEC3G | 0.49269214 | 2.00327 | < 0.001 | 0.006212312 |
| RECRUITMENT OF MITOTIC CENTROSOME PROTEINS AND COMPLEXES | 0.4652946 | 1.9718788 | < 0.001 | 0.0077209 |
| AXON GUIDANCE | 0.37230575 | 1.95611 | < 0.001 | 0.008599579 |
| SCF BETA TRCP MEDIATED DEGRADATION OF EMI1 | 0.484773 | 1.9473646 | < 0.001 | 0.009275142 |
| REGULATION OF ORNITHINE DECARBOXYLASE ODC | 0.4825054 | 1.9217154 | < 0.001 | 0.011184356 |
| MITOTIC M G1 PHASES | 0.38666242 | 1.9204922 | < 0.001 | 0.011307008 |
| SIGNALING BY ROBO RECEPTOR | 0.54419017 | 1.9131768 | < 0.001 | 0.011887343 |
| FORMATION OF TUBULIN FOLDING INTERMEDIATES BY CCT TRIC | 0.5788052 | 1.8959764 | < 0.001 | 0.0132828 |
| P130CAS LINKAGE TO MAPK SIGNALING FOR INTEGRINS | 0.63985723 | 1.8917065 | 0.002242153 | 0.013693307 |
| ENERGY DEPENDENT REGULATION OF MTOR BY LKB1 AMPK | 0.6105954 | 1.8908657 | < 0.001 | 0.013682265 |
| AUTODEGRADATION OF CDH1 BY CDH1 APC/C | 0.45841932 | 1.8844975 | < 0.001 | 0.014151104 |
| INHIBITION OF INSULIN SECRETION BY ADRENALINE NORADRENALINE | 0.5502511 | 1.8698757 | 0.004784689 | 0.01577267 |
| DESTABILIZATION OF MRNA BY AUF1 HNRNP D0 | 0.46294895 | 1.8583478 | < 0.001 | 0.017308583 |
| MITOTIC G2 G2 M PHASES | 0.42301303 | 1.8577948 | < 0.001 | 0.01730274 |
| REGULATION OF MITOTIC CELL CYCLE | 0.42331395 | 1.8523718 | < 0.001 | 0.017783675 |
| ER PHAGOSOME PATHWAY | 0.45423186 | 1.8411376 | < 0.001 | 0.0194142 |
| CDK MEDIATED PHOSPHORYLATION AND REMOVAL OF CDC6 | 0.46669722 | 1.8394731 | < 0.001 | 0.019573752 |
| G BETA GAMMA SIGNALLING THROUGH PI3KGAMMA | 0.5516966 | 1.8369927 | 0.002262444 | 0.019893283 |
| THROMBOXANE SIGNALLING THROUGH TP RECEPTOR | 0.5546398 | 1.8334056 | 0.009237875 | 0.02019652 |
| CROSS PRESENTATION OF SOLUBLE EXOGENOUS ANTIGENS ENDOSOMES | 0.46626863 | 1.8325262 | < 0.001 | 0.020241262 |
| REGULATION OF AMPK ACTIVITY VIA LKB1 | 0.6282577 | 1.8308858 | < 0.001 | 0.020418137 |
| HOST INTERACTIONS OF HIV FACTORS | 0.38340074 | 1.8304092 | < 0.001 | 0.020490842 |
| APC C CDC20 MEDIATED DEGRADATION OF MITOTIC PROTEINS | 0.43205938 | 1.8181952 | < 0.001 | 0.022108192 |
| SIGNAL AMPLIFICATION | 0.5096442 | 1.816911 | 0.002364066 | 0.022154026 |
| PROSTACYCLIN SIGNALLING THROUGH PROSTACYCLIN RECEPTOR | 0.5756845 | 1.805098 | 0.002398082 | 0.0237121 |
| CDT1 ASSOCIATION WITH THE CDC6 ORC ORIGIN COMPLEX | 0.4404016 | 1.8014396 | < 0.001 | 0.024302073 |
| AUTODEGRADATION OF THE E3 UBIQUITIN LIGASE COP1 | 0.45105973 | 1.7888427 | 0.002415459 | 0.026457857 |
| GLUCAGON SIGNALING IN METABOLIC REGULATION | 0.49052048 | 1.7840647 | < 0.001 | 0.026994882 |
| REGULATION OF INSULIN SECRETION BY GLUCAGON LIKE PEPTIDE1 | 0.46110502 | 1.7792808 | 0.004975124 | 0.027836313 |
| REGULATION OF APOPTOSIS | 0.43052307 | 1.7744424 | < 0.001 | 0.028887268 |
| DNA REPLICATION | 0.35073006 | 1.7696922 | < 0.001 | 0.029675506 |
| CELL CYCLE MITOTIC | 0.32204235 | 1.7489924 | < 0.001 | 0.032654043 |
| PLATELET ACTIVATION SIGNALING AND AGGREGATION | 0.33945197 | 1.7335683 | < 0.001 | 0.034900293 |
| POST CHAPERONIN TUBULIN FOLDING PATHWAY | 0.5681244 | 1.7331092 | 0.009090909 | 0.034972716 |
| ACTIVATION OF NF KAPPAB IN B CELLS | 0.4113944 | 1.7256137 | 0.002525253 | 0.03624478 |
| PROTEIN FOLDING | 0.4404603 | 1.7220641 | < 0.001 | 0.037190825 |
| MYOGENESIS | 0.49990174 | 1.7121415 | 0.011286682 | 0.039187293 |
| ADP SIGNALLING THROUGH P2RY12 | 0.5304115 | 1.7079254 | 0.00896861 | 0.039884042 |
| GRB2 SOS PROVIDES LINKAGE TO MAPK SIGNALING FOR INTERGRINS | 0.58723575 | 1.699356 | 0.006976744 | 0.04175493 |
| P53 INDEPENDENT G1 S DNA DAMAGE CHECKPOINT | 0.4222017 | 1.6951325 | 0.002352941 | 0.042560223 |
| INTEGRIN CELL SURFACE INTERACTIONS | 0.38336936 | 1.6892639 | < 0.001 | 0.043578427 |
| ADP SIGNALLING THROUGH P2RY1 | 0.50572884 | 1.6892458 | 0.009615385 | 0.043502424 |
| PYRIMIDINE METABOLISM | 0.51263106 | 1.680901 | 0.016241299 | 0.0443648 |
| L1CAM INTERACTIONS | 0.37919116 | 1.6799189 | 0.002638523 | 0.04459992 |
| PREFOLDIN MEDIATED TRANSFER OF<br>SUBSTRATE TO CCT TRIC | 0.4865133 | 1.6771183 | 0.00990099 | 0.04532867 |
| INTEGRATION OF ENERGY METABOLISM | 0.3586314 | 1.6719067 | < 0.001 | 0.046673577 |

**Supplemental table 2.**
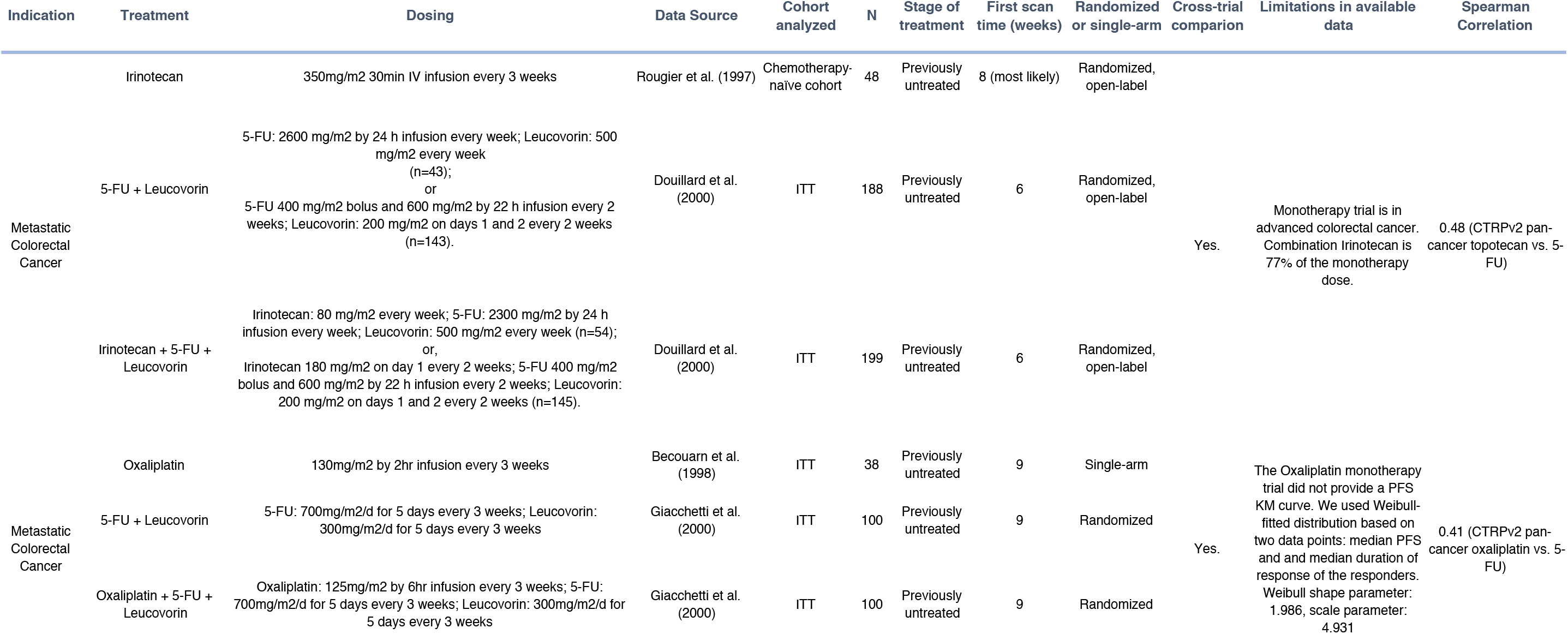
Drug additivity analysis for clinical 5-FU-based chemotherapy trial.

